# Macrocycle-stabilization of its interaction with 14-3-3 increases plasma membrane localization and activity of CFTR

**DOI:** 10.1101/2022.01.03.473871

**Authors:** Loes M. Stevers, Madita Wolter, Graeme W. Carlile, Dwight Macdonald, Luc Richard, Frank Gielkens, David Y. Thomas, Sai Kumar Chakka, Mark L. Peterson, Helmut Thomas, Luc Brunsveld, Christian Ottmann

## Abstract

Impaired activity of the chloride channel CFTR is the cause of cystic fibrosis. 14-3-3 proteins have been shown to stabilize CFTR and increase its biogenesis and activity. Here, we report the identification and mechanism of action of a macrocycle stabilizing the 14-3-3/CFTR complex, a first-in-class molecular glue. This molecule rescues plasma membrane localization and chloride transport of F508del-CFTR and works additively with the CFTR pharmacological chaperone corrector lumacaftor (VX-809).

---

The CFTR protein is a cyclic adenosine 5⍰-monophosphate (cAMP)–regulated transporter with anion channel activity that conducts Cl^−^ on the apical surface of bronchial epithelial cells. The F508del-CFTR mutation is the most frequent cause of cystic fibrosis (CF)^1^. It codes for a mutant protein that is recognized as misfolded and retained in the endoplasmic reticulum but, if induced to traffic to the plasma membrane, it is almost fully functional^2^. Cell-based assays have identified compounds that facilitate the trafficking of F508del-CFTR to the plasma membrane. So far, the most effective compounds for F508del-CFTR have been pharmacological chaperones that bind to the mutant CFTR molecule and assist its correct folding^1^. To amplify the function of the corrected F508del-CFTR, the potentiator ivacaftor, (Kalydeco®) that corrects the G551D (non-trafficking mutation) is included in combination with pharmacological chaperones^3^. There have been two approvals of such drug combinations that correct the trafficking of the F508del-CFTR, ivacaftor/lumacaftor (Orkambi®)^4^ in 2015 and ivacaftor/tezacaftor (Symdeko®)^5^ in 2018. More recently (2019), the triple combination ivacaftor/tezacaftor/elexacaftor (Trikafta®) has been approved^6^. The need for these combinations is that single molecules give low levels of correction and, although a new corrector VX-445 (elexacaftor) has been recently described that appears to give clinically significant levels of correction over a range of CFTR mutations, optimal results still require combination therapies^7,8^.

These molecules are believed to target CFTR directly and function as molecular chaperones. In particular, ivacaftor and an investigational drug from Galapagos (GLPG1837) have been shown to bind CFTR at the protein/plasma membrane interface with half of the molecules’ surfaces exposed to the lipid bilayer^9^. Although the above-described drugs represent breakthrough therapies for cystic fibrosis, it is still highly informative to explore further aspects of CFTR biology for modulation by small molecules to potentially improve clinical outcomes. For example, protein-protein interactions (PPIs) have been the focus of drug discovery and chemical biology for some time^10–13^, triggered by the seminal examples of the natural products rapamycin and FK506^14^, the approval of venetoclax (Venclexta®) as the first Bcl-2 inhibitor^15^, and the tremendous clinical and economic success of lenalidomide (Revlimid®)^16^. In 2015, Pankow et al. reported a CFTR interactome analysis of both wildtype (wt) and F508del-CFTR^17^. In this study, 638 partner proteins of CFTR were identified and an extensive remodeling of the F508-del interactome upon rescue of mutated CFTR function by low temperature (26 - 30°C.) or HDAC inhibition was demonstrated. Furthermore, RNA interference was used to identify proteins whose specific knockdowns rescue or reduce CFTR function in the F508-del mutant, many of which are involved in the degradation machinery, quality control, or membrane trafficking processes.^17^ These findings suggest that modulation of the CFTR interactome could contribute to a therapeutically beneficial restoration of impaired CFTR function. Interestingly, among the proteins that in this study were consistently found in the CFTR interactome of both wt and F508-del, were the 14-3-3 proteins, important regulators of Ser/Thr-phosphorylated proteins.

14-3-3 proteins are dimeric proteins that have been shown to bind to the disordered regulatory (R) domain of CFTR, facilitate trafficking to the plasma membrane and enhance ion channel activity^18,19^. We have previously reported the crystal structure of 14-3-3 in complex with a number of phosphopeptides derived from CFTR and have shown that the natural product fusicoccin A (FC-A) can stabilize the 14-3-3/CFTR interaction and promote plasma membrane localization of F508del-CFTR^20^. Since the structural complexity of fusicoccanes poses a significant challenge for medicinal chemistry optimization, we sought to identify novel synthetic chemotypes that are able to stabilize the 14-3-3/CFTR complex. To this end, we screened 5760 compounds of Cyclenium’s proprietary small-molecule macrocycle library employing a fluorescence polarization (FP) assay for the binding of the di-phosphorylated CFTR-derived synthetic peptide CFTRpS753pS768 to 14-3-3β (SupplementaryFig. 1a). In the single concentration screen, 24 hits were identified (Supplementary Fig. 1b) of which seven (7) showed concentration-dependent stabilization (Supplementary Fig. 1c). These compounds can be grouped into four (4) chemotypes (Supplementary Fig. 2) which prompted us to design and synthesize a second, focused validation library of 480 macrocycles. FP screening of this library added another eight (8) compounds (Supplementary Fig. 3a) as validated stabilizers of the 14-3-3β/CFTRpS753pS768 interaction to the initial set (Supplementary Fig. 3b). In the presence of these compounds, the apparent K_d_ for the CFTR peptide binding to 14-3-3 was increased significantly (Supplementary Fig. 4).

In order to gain structural information on the PPI stabilizing activity of these compounds, co-crystallization trials of 14-3-3β in complex with eight (8) of the validated hits were performed. Diffraction-quality crystals could be optimized for the complex with CY007424 (Fig. 1a) and the structure solved at 1.78 Å. The 14-3-3β/CFTRpS753pS768/CY007424 complex crystallized as a tetramer in the asymmetric unit with two 14-3-3 dimers, two copies of the CFTR peptide and two molecules of CY007424 (Supplementary Fig. 5) with electron density covering the entire macrocyclic molecule (Fig. 1b). Of the 28 residues of the CFTRpS753pS768 peptide, 21 amino acids could be built into the model (Fig. 1c). CY007424 binds close to pS753 of the CFTR peptide and establishes contacts to 14-3-3, as well as to the peptide (Fig. 1c, Supplementary Fig. 6). The tyrosine moiety of CY007424 is embedded in a shallow hydrophobic cleft formed by Pro750 and Ile752 of CFTR and Leu229 of 14-3-3β (Fig. 1c, Supplementary Fig. 6a). The two main ring thioether phenyls are engaged in a hydrophobic interaction with the hydrocarbon part of Arg58, Ser59 and Arg62 of 14-3-3 (Fig. 1c, Supplementary Fig. 6b). There is a polar interaction between the main chain nitrogen and carbonyl oxygen of Arg751 of CFTR and the corresponding ring nitrogen and carbonyl of the tyrosine moiety of CY007424 (Supplementary Fig. 6c). A third polar contact is established between the arginine moiety of CY007424 and Asn52 of 14-3-3β (Supplementary Fig. 6d).

**Fig. 1.**
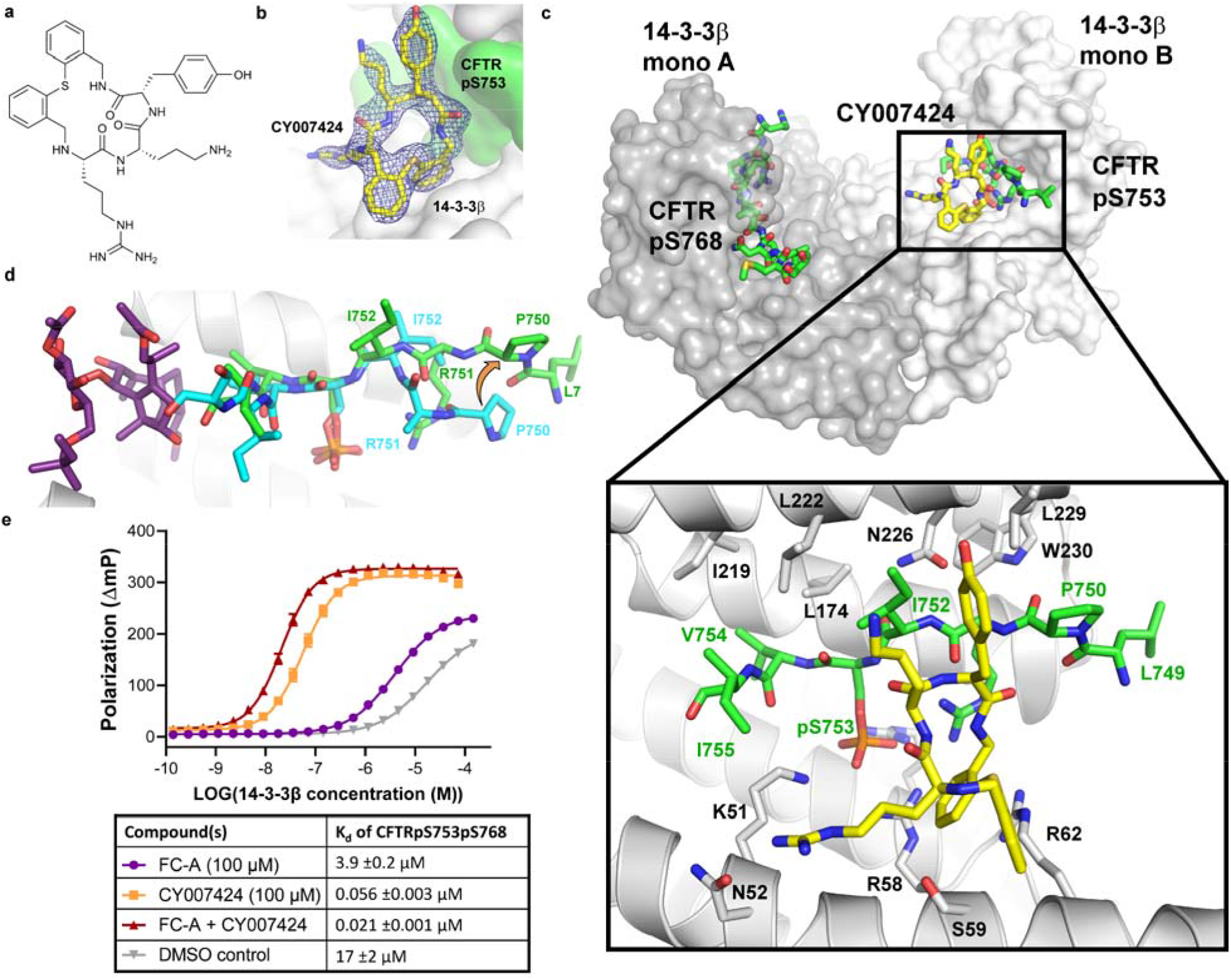
Macrocycle stabilization of the 14-3-3/CFTR interaction. **a**, Chemical structure of CY0072424. **b**, Electron density map (2Fo-Fc, 1σ) of CY007424 bound to the interface of 14-3-3β and CFTRpS753. **c**, Crystal structure of the 14-3-3β homodimer complexed with CFTRpS753pS768 and CY007424. **d**, Overlay of the CFTRpS753 peptide motif bound to 14-3-3 in the presence of FC-A (cyan sticks) or CY007424 (green sticks). **e**, FP assay of FITC-labeled CFTRpS753pS768 (10 nM) with 14-3-3β in the presence of 100 μM of FC-A, CY007424, or both compounds.

Comparison of the binding mode of the previously established FC-stabilized CFTRpS753 interaction motif with the current CY007424-stabilized structure revealed a considerable conformational change induced by CY007424 in the N-terminus of the peptide (Fig. 1d). The positions of P750 and R751 flip by around 180° and the side chains of L749 and R751 become visible in the electron density. In the case of R751, this allows a polar interaction between the phosphate of pS753 and the terminal amino group of this arginine, as well as a direct contact with CY007424 (Supplementary Fig. 6d). The observed conformational changes are necessary to establish the above-described binding mode, especially the accommodation of the tyrosine moiety. This means that CY007424 either “selected” the observed peptide conformation from an ensemble of states that this flexible peptide can adopt or an “induced-fit” adaptation of the peptide took place in the presence of the macrocycle.

In accordance with the different binding sites of the two 14-3-3/CFTR stabilizers FC-A and CY007424, simultaneous treatment resulted in an additive effect and increased the apparent affinity of the CFTR peptide to 14-3-3 by three orders of magnitude, from 17 μM to 21 nM (Fig. 1e). CY007424 is a much more potent compound than FC-A^20^, stabilizing the complex by a factor of more than 300x, whereas FC-A shows an about 4.5x stabilization at a concentration of 100 μM (Fig. 1e).

Twelve (12) validated hit compounds from the initial screening and focused validation library were tested for their effect on F508del-CFTR trafficking. Baby hamster kidney (BHK) cells expressing 3HA-tagged F508del-CFTR were treated for 24 hours with 10μM compound. After fixation of the cells, a combination of mouse monoclonal anti-HA antibody and anti-mouse IgG conjugated with FITC was used to detect F508del-CFTR that had trafficked to the plasma membrane as described previously^21^. CY007424 showed the strongest increase of F508del-CFTR trafficking towards the plasma membrane, followed by CY007491, CY007476, and CY007288 (Fig. 2a). Additionally, a fluorescence imaging plate reader (FLIPR) Membrane Potential (FMP) assay was performed. This assay measures real-time membrane potential changes associated with ion channel activation and ion transporter proteins. F508del-CFTR expressing BHK cells were incubated for 24 hours with 20μM compound before treatment with the potentiator genistein, FMP dye, and activator forskolin. The fluorescence intensity is thus a measure for the function of the CFTR protein in the plasma membrane of these cells. The F508del-CFTR corrector VX-809 (lumacaftor) was used as a positive control in this assay. CY007424 has a clear corrector function on F508del-CFTR, while the other macrocycles show no significant increase in CFTR function compared to the DMSO control (Fig. 2b). When the cells were incubated with the CFTR inhibitor 172 (INH172) this effect was neutralized (Fig. 2c). Interestingly, the combination of VX-809 and CY007424 shows an additive effect on CFTR function in the cell membrane (Fig. 2c). Another cellular assay was performed with the Ussing Chamber, which detects and quantifies transport of ions across epithelial tissue^22^. F508del-CFTR expressing cystic fibrosis epithelial (CFBE) cells were treated for 18 hours with 20μM CY007424. After forskolin and genistein stimulation, the short-circuit current of the cells was measured. The CY007424-treated CFBE cells showed a higher conductance than the DMSO control treated cells. Also here, the combination of CY007424 and VX-809 showed an additive response of 150% compared to VX-809 alone (Fig. 2d).

**Fig. 2.**
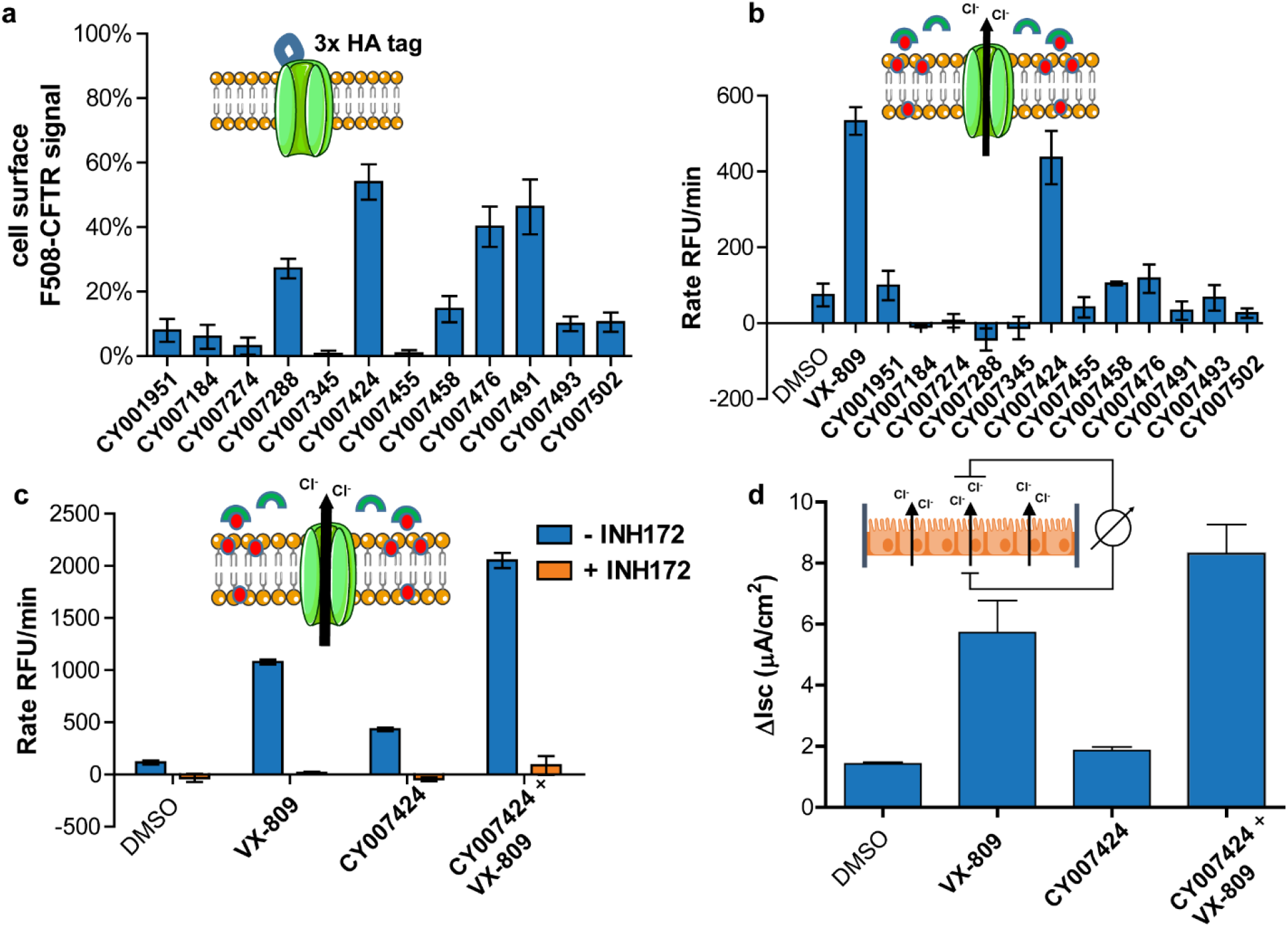
Cellular activity of macrocycles stabilizing the 14-3-3/CFTR PPI. **a**, Surface expression of 3HA-tagged F508del-CFTR in BHK cells incubated for 24 h with 10 μM compound and treated with mouse monoclonal anti-HA antibody and anti-mouse IgG conjugated with FITC. **b, c**, FLIPR Membrane Potential (FMP) assay of F508del-CFTR expressing BHK cells treated for 24 h with the macrocycles (b) or with CY007424 and/or VX-809, with and without CFTR inhibitor INH172 (c). **d**, Ussing chamber experiment of F508del-CFTR expressing CFBE cells treated for 18 h with CY007424 or VX-809 or a combination of both. CFTR activity is assayed by measurement of short-circuit current of cells after stimulation by forskolin and the potentiator genistein.

In conclusion, the small molecule macrocycles presented in this study are the first synthetic compounds that have been shown to stabilize the interaction of the CF-related chloride channel CFTR with 14-3-3 proteins in an unprecedented mode of action. Since 14-3-3 proteins are positive regulators of CFTR that facilitate forward trafficking to the plasma membrane and stabilize the functional fold of the channel, these compounds are useful tools to evaluate the CFTR/14-3-3 interaction as a potential target for therapeutic intervention in CF.

## Online content

Any methods, additional references, Nature Research reporting summaries, source data, statements of code and data availability and associated accession codes are available at….

## Acknowledgements

JWH, GWC, and DYT are supported by grants from the Canadian Institutes of Health Research and Cystic Fibrosis Canada. DYT is the Canada Research Chair in Molecular Genetics. This work is funded by the H2020 Marie Curie Actions of the European Commission through the TASPPI project, grant Agreement 675179 and by the Netherlands Organization for Scientific Research via Gravity Program 024.001.035.

## Additional information

Supplementary Information is available for this paper at

Reprints and permissions information is available at www.nature.com/reprints.

Correspondence and requests for materials should be addressed to

## Methods

### Reagents

Fusicoccin A (FC-A) was obtained from Enzo Life Sciences BVBA. All small molecule macrocycle compounds and libraries were synthesized as described in the Supplementary Information and provided by Cyclenium Pharma.

### Peptide synthesis

The CFTR peptides were synthesized as previously described.^20^

### Expression of 14-3-3

His_6_-tagged 14-3-3 isoforms (full-length and ΔC) were expressed in NiCo21(DE3) competent cells (0.4 mM IPTG, overnight at 18 °C), with a pPROEX HTb plasmid, and purified with a nickel column. The His6-tag was cleaved-off with TEV-protease and a second purification was done by size exclusion chromatography. The proteins were dialyzed against FP, ICT, or crystallization buffers before usage (described below).

### Fluorescence polarization (FP) assay

The FITC-labeled peptides were dissolved in FP buffer (10 mM HEPES pH 7.4, 150 mM NaCl, 0.1% Tween20, 1 mg/mL BSA) to a final concentration of 100 nM or 10 nM. For the dose-response assays, a 14--3-3β concentration of 10 μM was used while titrating the compounds in a two-times dilution series and, for the other assays, a dilution series of 14-3-3β was made with constant concentration of compound. The dilution series were made in 384 Corning Black Round Bottom well plates and their polarization was measured with a Tecan Infinite F500 plate reader (ex. = 485 nm, em. = 535 nm).

### HTS FP assay

The HTS FP assay was set up in 384 Corning Black Round Bottom well plates. Each well contained 20 μL sample with 10 μM 14-3-3β and 100 nM FITC-labeled CFTRpS753pS768 peptide dissolved in FP buffer (10 mM HEPES pH 7.4, 150 mM NaCl, 0.1% Tween20, 1 mg/mL BSA). Compound was added using a pin tool system adding approximately 0.1 μL to each well, dependent on the viscosity of the solution. Each plate contained 16 wells with negative controls (DMSO) and 16 wells with positive control (FC-A). The plates were incubated for 30 min in the dark at RT before measuring the polarization using a PHERAstar FS plate reader (ex. = 485 nm, em. = 520 nm).

### Crystallography

The 14-3-3β protein was C-terminally truncated after T232 to improve crystallization. For crystallization, the 14-3-3βΔC/CFTR_pS753pS768/CY07424 complex was mixed in a 2:1.5:4 molar stoichiometry with a final protein concentration of 15 mg/mL in crystallization buffer (25 mM HEPES, 0.1 M NaCl, 2 mM DTT, pH 7.4). This was set up for hanging-drop crystallization in a 1:1 ratio with Qiagen Cryos Suite #44 crystallization liquor (0.09 M HEPES sodium salt, pH 7.5, 1.26 M tri-Sodium citrate, 10% (v/v) Glycerol) with an extra 2% of Glycerol (SigmaAldrich). Crystals were fished out after 2 weeks of incubation at 4°C and flash-cooled in liquid nitrogen. Diffraction data was collected at the PETRA III P11 beamline (DESY, Hamburg, Germany). The dataset was indexed and integrated using XDS^23^ and scaled using SCALA^24^. The structure was phased by molecular replacement, using PDB ID 2C23^25^ as search model, in Phaser^26^, Coot^27^ and phenix.refine^28^ were used in alternating cycles of model building and refinement. See Table S1 for data collection, structure determination, and refinement.

### CFTR trafficking assay

The trafficking assay was performed as previously described by Carlile *et al*. ^29^ In brief, 3HA-tagged F508del-CFTR expressing baby hamster kidney (BHK) cells were seeded in 96-well plates (Corning half area, black-sided, clear bottom) at 15,000 cells per well. After 24 h incubation at 37°C, the cells were treated with 10 μM of compound for 24 h (final DMSO concentration 1% v/v). The cells were fixated with 4% paraformaldehyde, washed with PBS, and then blocked with fetal bovine serum (5% in PBS). Mouse monoclonal anti-HA antibody (Sigma, 1:150 dilution in PBS) was incubated overnight, and, after three washes with PBS, the background fluorescence was measured in a plate reader (488 nm excitation, 510 nm emission). The secondary antibody anti-mouse IgG conjugated with FITC (Sigma, 1:100 dilution in PBS) was incubated for 1 h, the cells were washed three times with PBS and analyzed in the plate reader again. Background fluorescence was subtracted from the signal, after which the signal was normalized to the DMSO control and wt-CFTR expressing cells.

### FLIPR Membrane Potential assay

The FLuorescence Imaging Plate Reader (FLIPR) Membrane Potential (FMP) assay is based on the technique developed by Van Goor *et al*.^30^ F508del-CFTR expressing baby hamster kidney (BHK) cells were incubated with the compounds for 24 h at 37°C. The growth medium was removed from the cells by inverting the plate and FMP dye (Molecular Devices, Part #R8042) including the potentiator genistein (Sigma, G6649) was added back in 70 ul of low Cl^-^ containing buffer (160 mM NaGluconate, 4.5 mM KCl, 2 mM CaCl_2_, 1 mM MgCl_2_, 10 mM D-Glucose, 10 mM HEPES (pH7.4)). The plates were incubated for 5 min at RT before activation of CFTR in the plate reader (SynergyMX) with the addition of 14 µL of FMP dye in low Cl^-^ buffer containing 6x forskolin (Sigma, F6886) following a 2 min baseline read. Fluorescence intensity was monitored for 5 min following CFTR activation. Reported is the rate of fluorescence intensity change over time.

### Ussing Chamber assay

Primary F508del-CFTR expressing CFBE cells were grown in air-liquid interface culture conditions until fully differentiated. The cells were treated for 18 h with 20 μM CY07424, 1 μM VX-809 or the combination of the two compounds. Once the cells were mounted in the Ussing Chambers, 10 μM of forskolin, the potentiator genistein and CFTR inhibitor INH172 was added in the chambers before measuring the short-circuit current over the CFBE cells as described by Hug *et al*.^31^

