## Supporting Information for "Macrocycle-stabilization of its interaction with 14-3-3 increases plasma membrane localization and activity of CFTR"

### Supplementary Information

Supplementary Information contents:

SUPPLEMENTARY Figures

SUPPLEMENTARY Tables

SUPPLEMENTARY Note: Compound Synthesis and Characterization

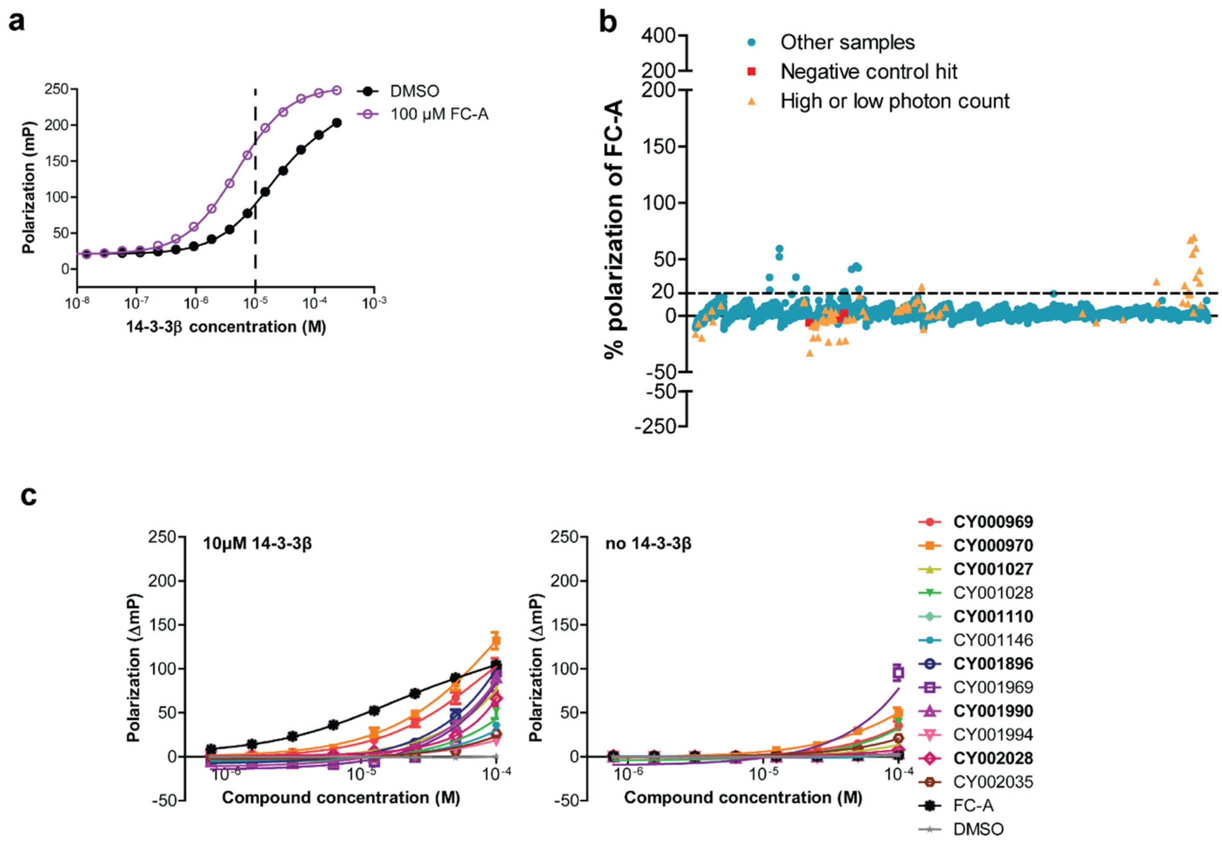

**Fig. S1 | Screening for stabilizers of the 14-3-3β/CFTRpS753pS768 complex. a.** Fluorescence polarization (FP) assay of fluorescein isothiocyanate (FITC)-labeled CFTRpS753pS768 peptide (100 nM) with 14-3-3β, in the absence (black) or presence (purple) of 100 µM fusicoccin A (FC-A). The 14-3-3β concentration used for the high throughput screening (HTS) FP assay was 10 µM (dashed line). Background polarization was subtracted from all values. Mean of three experiments, standard deviation (SD) error bars are smaller than the data point symbols. **b.** HTS FP assay of the Cyclenium library. The samples contain 100 nM of FITC-CFTR_pS753pS768 peptide, 10 µM 14-3-3β, and approximately 50 µM compound, dependent on the viscosity of the solution. FC-A (100 µM) was added as a positive control and DMSO as the negative control. Hit compounds are defined as a polarization increase which is higher than 20% of the response of FC-A. Red squares are compounds that appeared positive in the negative control without 14-3-3. Orange triangles gave a suspicious high or low total photon count and need to be treated with care. **c.** The eight-point dose-response FP follow-up assay of the Cyclenium library hit compounds stabilizing the interaction between 14-3-3 (10 µM) and labeled CFTRpS753pS768 (100 nM) peptide. Background polarization was subtracted from all values. Mean of three experiments, error bars are SD. Compounds selected for further analysis are displayed in bold.

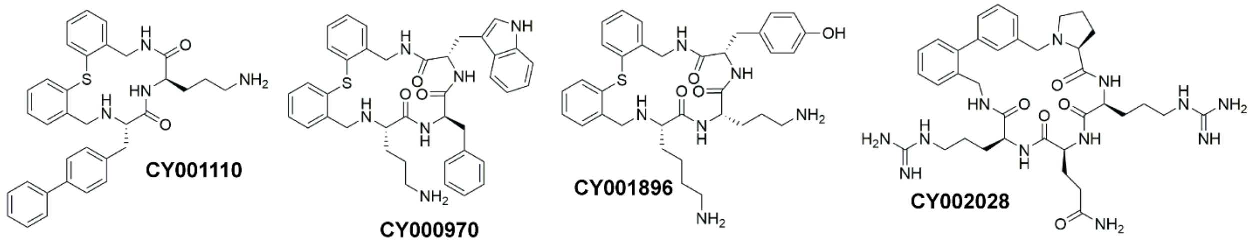

**Fig. S2 | Representatives from the four chemotypes of active compounds after screening for stabilizers of the 14-3-3β/CFTRpS753pS768 complex.**

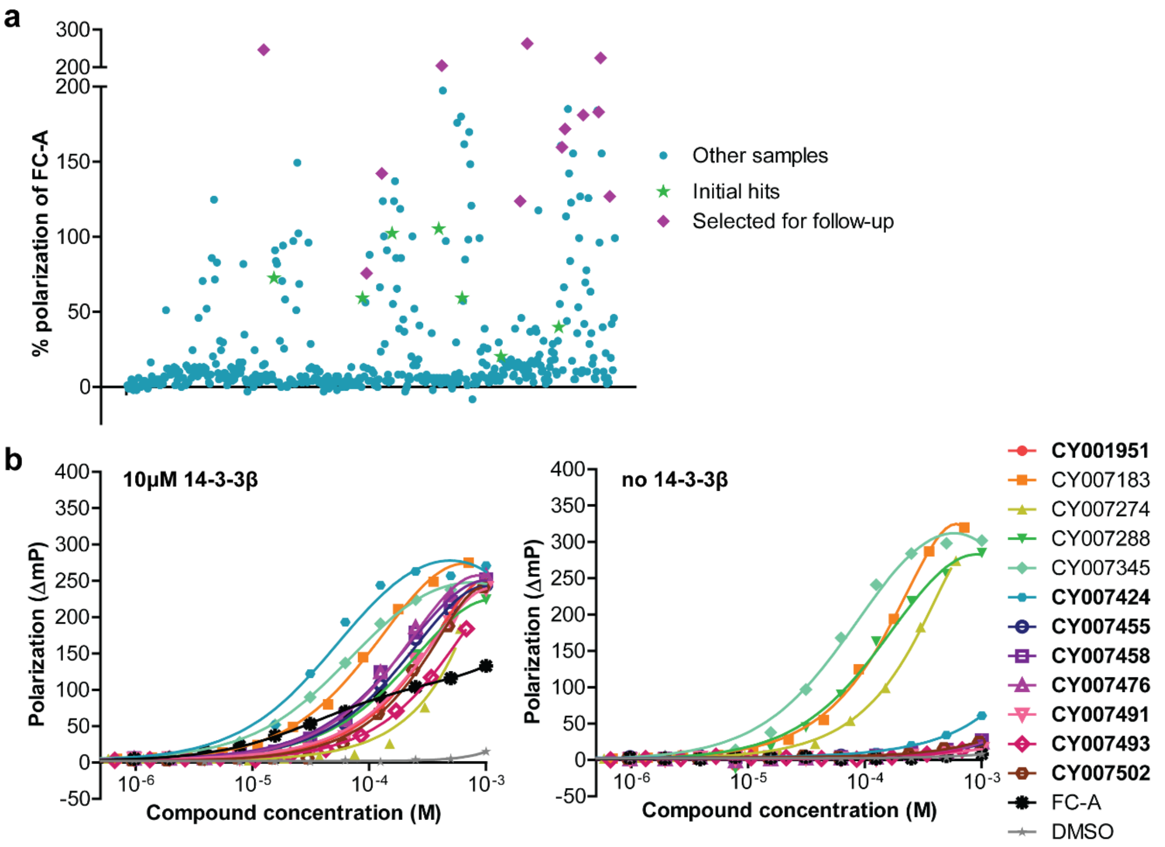

**Fig. S3 | Screening results of the hit validation library that consisted of resynthesized primary hit compounds and newly designed macrocycles to explore structure-activity relationships. a.** HTS FP of all 480 compounds. The samples contain 100 nM of FITC-CFTRpS753pS768 peptide, 10 µM 14-3-3β, and approximately 125 µM compound, dependent on the compound stock concentration. The positive control was FC-A (100 µM) and negative control was DMSO. Green stars are the initial hit compounds from previous library and the purple diamonds are selected for the follow-up dose-response assay. **b.** Dose-response FP follow-up assay of the selected hit compounds stabilizing the interaction between 14-3-3β (10 µM) and labeled CFTRpS753pS768 (100 nM) peptide. Background polarization was subtracted from all values. Compounds selected for further analysis are displayed in bold.

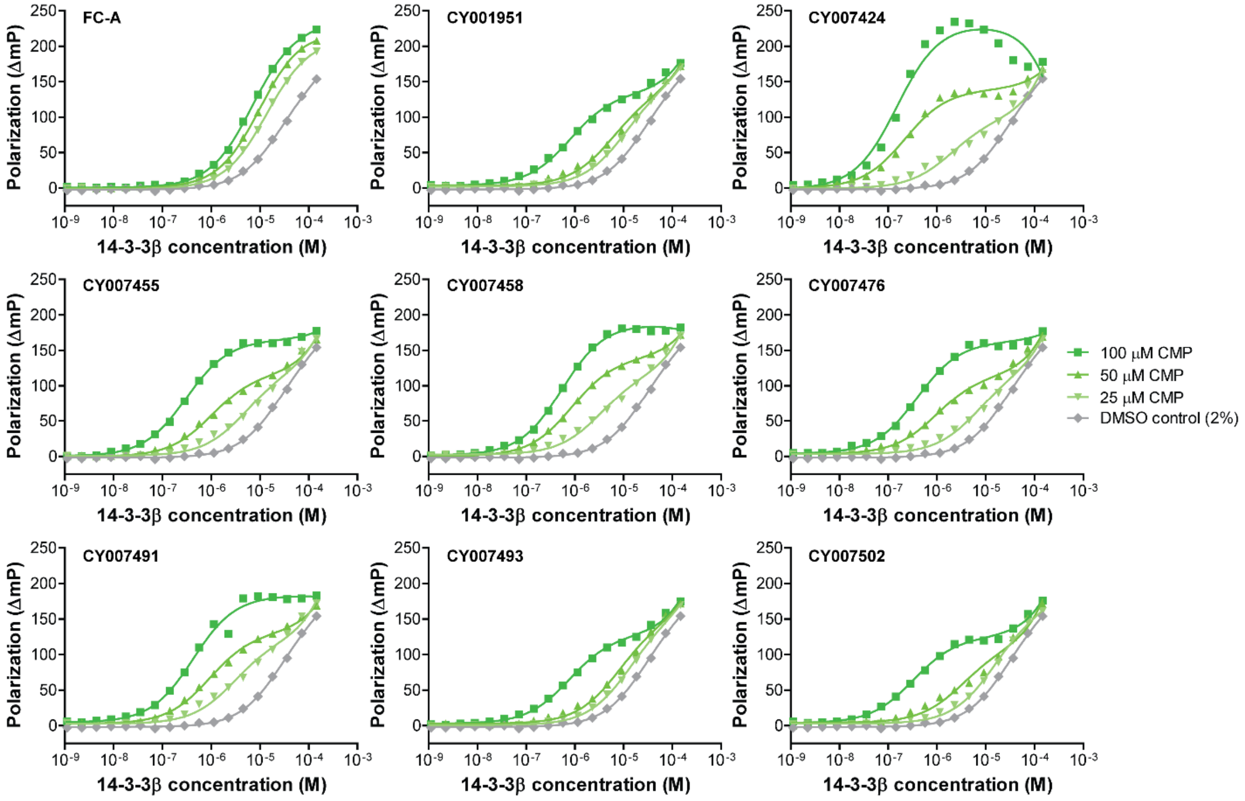

**Fig. S4 | Stabilization of the interaction between 14-3-3β and CFTRpS753pS768 by compounds from the hit validation library.** FP assay of FITC-labeled CFTRpS753pS768 with 14-3-3β in the presence of different concentrations of the compound.

**Table S1** Crystallographic statistics

| | 14-3-3βΔC / CFTRpS753pS768 / CY007424 |
|---|---|
| **Data collection** | |
| Wavelength (Å) | 1.03320 |
| Resolution (Å)[a] | 84.73 – 1.76 (1.79 – 1.76) |
| Space group | P 2 21 21 |
| Cell parameters (Å) | a = 70.37, b = 111.46, c = 130.41 |
| $CC_{1/2}$ (%)[a,b] | 0.999 (0.796) |
| $R_{merge}$ (%)[a,c] | 0.068 (1.095) |
| $R_{meas}$ (%)[a,c] | 0.074 (1.194 |
| Average $I/\sigma_{(I)}$[a] | 20.0 (2.1) |
| Completeness (%)[a] | 100.0 (100.0) |
| No. of unique reflections[a] | 102227 (4934) |
| Redundancy[a] | 11.8 (12.2) |
| Wilson B-factor ($Å^2$) | 9.24 |
| Mosaicity (°) | 0.62 |
| **Refinement** | |
| Number of non-solvent / solvent atoms | 7828 / 542 |
| $R_{work}$ / $R_{free}$ (%) | 19.6 / 23.9 |
| No. of reflections in the 'free' set | 5010 |
| R.m.s. deviations from ideal values | |
| bond lengths (Å) / bond angles (°) | 0.011 / 1.035 |
| Average B-factor (Å) | 28.93 |
| Ramachandran plot: favored / outlier residues (%) | 98.08 / 0.00 |
| Molprobity validation score | 2.40 |

[a] number in parentheses is for the highest resolution shell

[b] $CC_{1/2}$ = Pearson's intra-dataset correlation coefficient, reported by SCALA (version 3.3.22)

[c] Reported by SCALA (version 3.3.22)

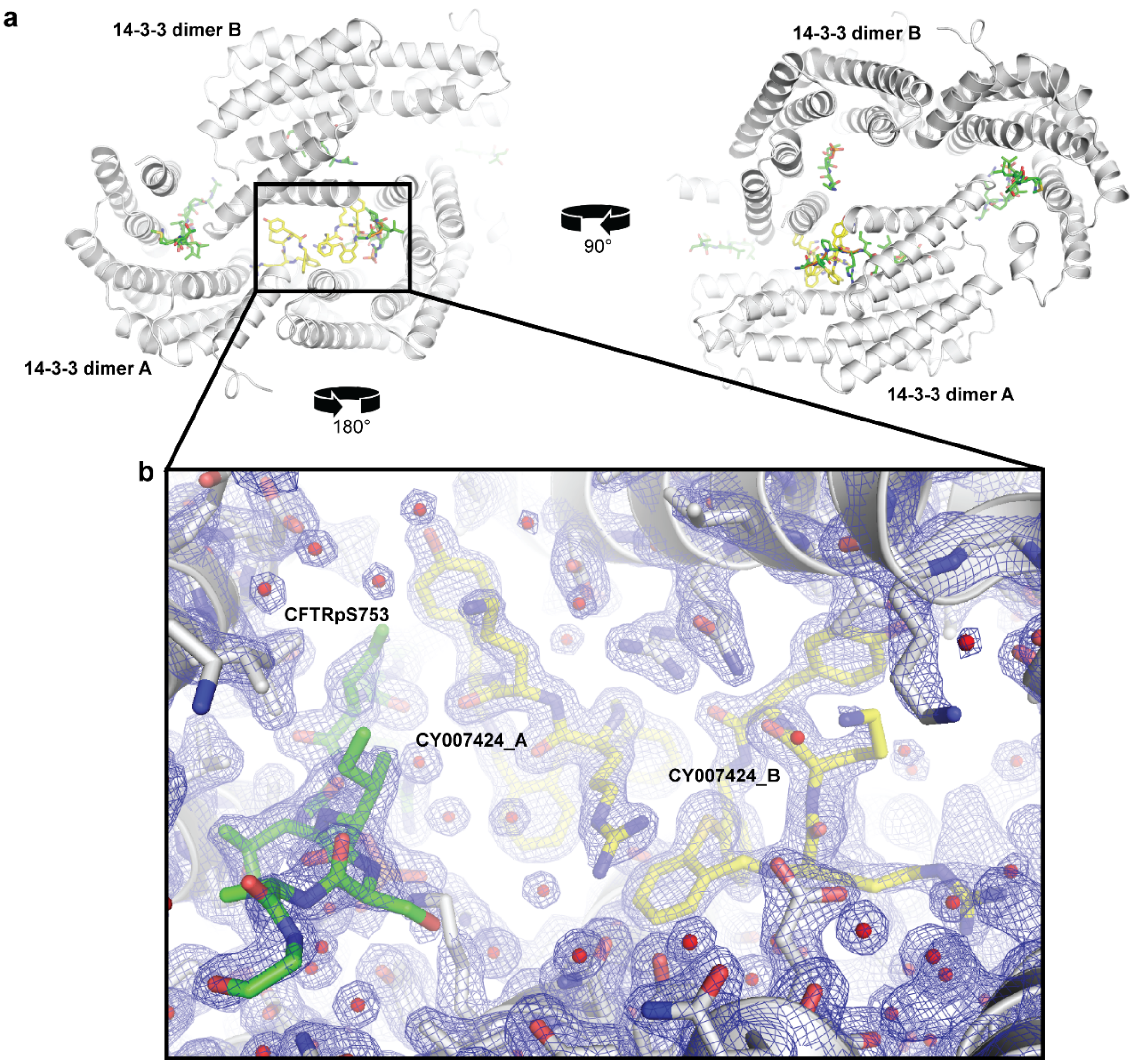

**Fig. S5 | The 14-3-3β/CFTRpS753pS768/CY007424 crystalizes as a tetramer in the asymmetric unit. a.** Two views of the tetramer consisting of two 14-3-3 dimers (A and B, white cartoon), two copies of the CFTRpS753pS768 peptide (green sticks), and two copies of CY007424 (yellow sticks). **b.** Detailed view of the final 2Fo-Fc electron density map highlighting the region of the two copies of CY007424 facilitating crystallization between the two dimers of 14-3-3.

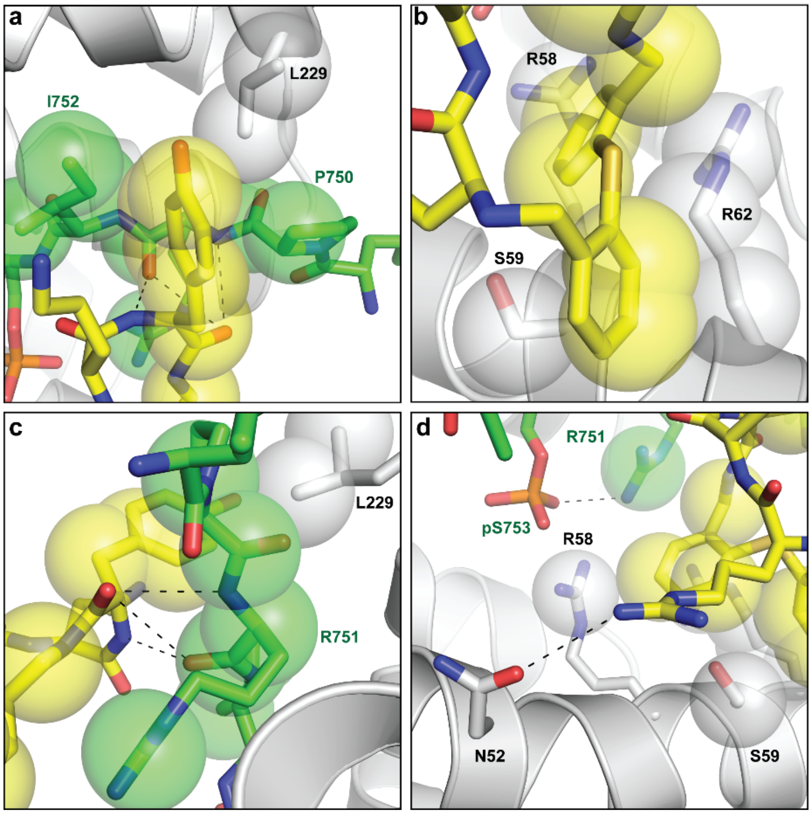

**Fig. S6 | Detailed view of the interactions of CY007424 binding to the complex of 14-3-3β and CFTRpS753pS768.** **a.** Interaction of the tyrosine moiety of CY007424 (yellow sticks and semi-transparent spheres) with P750 to I752 of the CFTRpS753pS768 peptide and L229 of 14-3-3β. **b.** Binding of the thio-bis-phenyl part of the core ring of CY007424 to the hydrocarbon part of R58, S59, and R62 of 14-3-3β. **c.** Polar interactions of the main-chain nitrogens and carbonyls of the tyrosine moiety of CY007424 and R751 of CFTRpS753pS768. **d.** Polar contact of the guanidinium group of CY007424 with N52 of 14-3-3β. In the presence of CY007424, the sidechain of R751 of CTRFpS753pS768 becomes visible and establishes a polar contact with the phosphate of pS753.

**SUPPLEMENTARY NOTE: Compound Synthesis and Characterization**

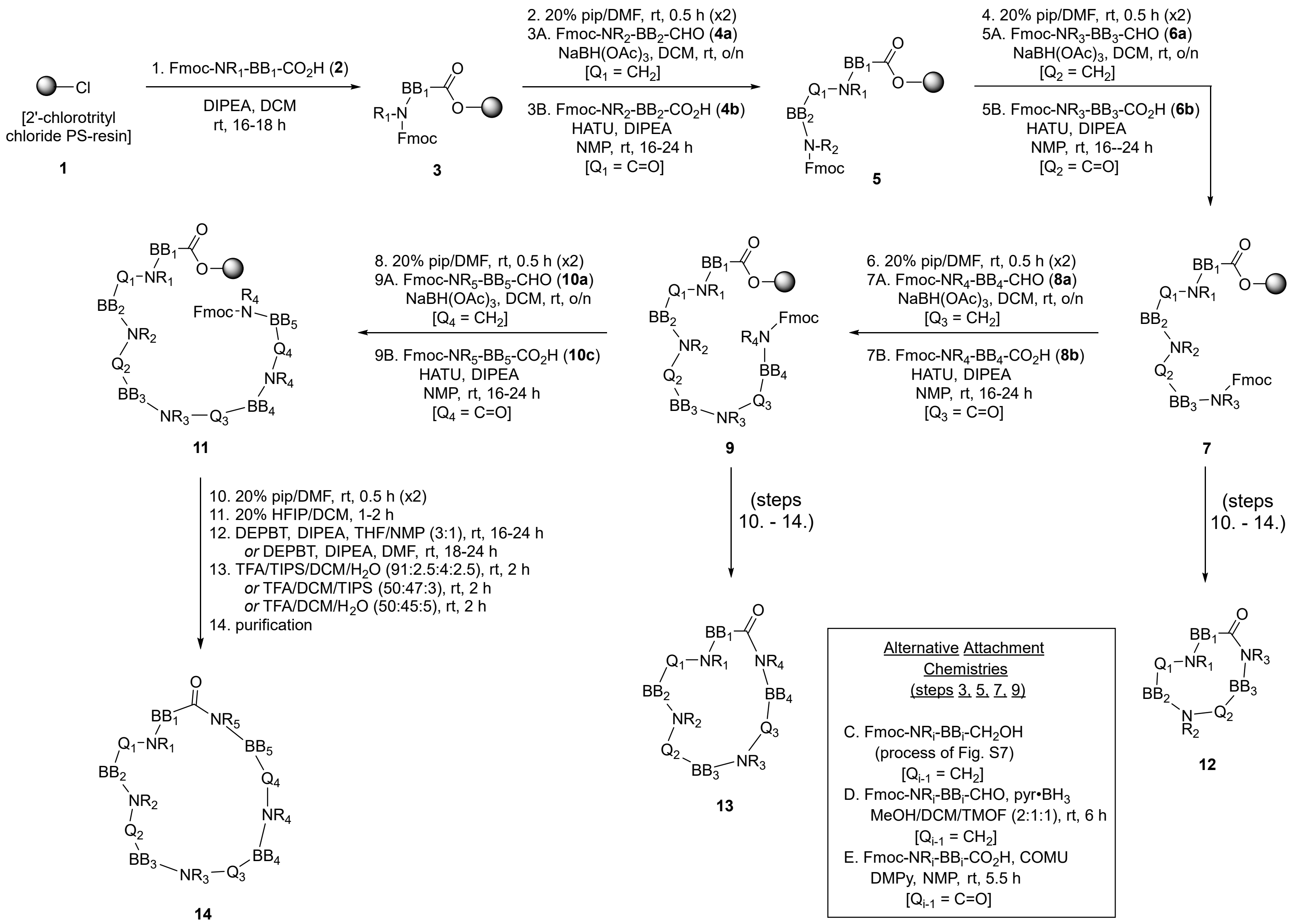

**Fig. S6 | General Synthetic Routes to Macrocyclic Compounds and Libraries**

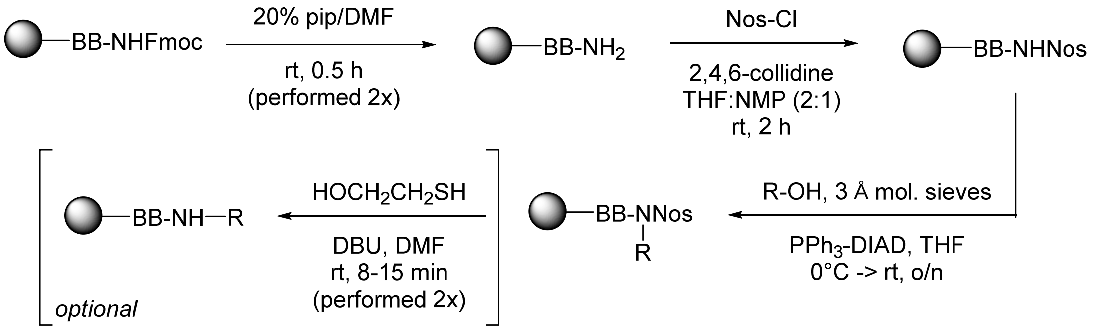

**Fig. S7 | General Process for Building Block Attachment using Fukuyama-Mitsunobu Reaction**

### Synthesis Procedures

#### General Information

Reagents and solvents were of reagent quality or better and were used as obtained from various commercial suppliers unless otherwise noted. Reaction solvents, such as DMF, DCM, DME and THF, were of DriSolv®, OmniSolv® (EMD Millipore, Darmstadt, Germany), or an equivalent synthesis grade quality. Reagent grade solvents were employed for: (i) deprotection, (ii) resin capping reactions and (iii) washing sequences. NMP used for coupling reactions was of analytical grade. DMF was adequately degassed by placing under vacuum for a minimum of 30 min prior to use. Ether refers to diethyl ether. Amino acids, Boc-, Fmoc- and Alloc-protected and side chain-protected derivatives, including those of N-methyl and unnatural amino acids, were obtained from commercial suppliers, including AAPPTec (Louisville, KY, USA), Advanced ChemTech (part of CreoSalus, Louisville, KY, USA), AstaTech (Bristol, PA, USA), Bachem (Bubendorf, Switzerland), Chem-Impex International (Wood Dale, IL, USA), Iris Biotech (Marktredwitz, Germany), Matrix Scientific (Columbia, SC, USA). Non-amino acid components were purchased from traditional vendors, such as Lancaster Synthesis (Ward Hill, MA, USA, subsidiary of Alfa Aesar, a Johnson Matthey Company), MilliporeSigma (St. Louis, MO, USA, subsidiary of Merck KGaA), and TCI America (Portland, OR, USA), or prepared utilizing literature methods. When N-protection of purchased material was required, this was accomplished through standard methodologies. Resins for solid phase synthesis were obtained from specialty suppliers, including AAPTech, Novabiochem (Oakville, ON, Canada, subsidiary of MilliporeSigma) and Rapp Polymere (Tübingen, Germany). Analytical TLC was performed on pre-coated plates of silica gel 60F254 (0.25 mm thickness) containing a fluorescent indicator.

$^{1}$H NMR spectra were recorded on a Bruker 500 MHz spectrometer equipped with a Double Resonance Broadband Probe (BBI) probe or a Bruker AVANCE II 700 MHz spectrometer equipped with a cryoprobe (5 mm CPDCH 13C-1H/D) and are referenced internally with respect to the residual proton signals of the indicated deuterated solvent. $^{13}$C NMR spectra also were recorded on the AVANCE II, at 176 MHz. Well-defined coupling patterns for observed resonances are indicated with standard abbreviations (s: singlet, d: doublet, t; triplet, q: quartet, br: broad, dd: doublet of doublets, dq: doublet of quartets, etc.) and coupling constants (*J*) with the number of protons represented by integration of each resonance denoted by "xH".

High resolution mass spectra (HRMS) for accurate mass measurements were performed on an Agilent Technologies LC-TOF 6224 instrument with either electrospray ionization (ESI) or atmospheric pressure chemical ionization (APCI). Aliquots of 0.2 uL were injected into the mass spectrometer using a 0.5 mL/min flow of 50% MeOH/50% $H_2O$ (containing 0.1% formic acid) mixture. The capillary voltage was set at 3000 V and mass spectra were acquired in the range 100-3000 *m/z*.

HPLC analyses were performed on an Agilent 1100 system equipped with a photodiode array (PDA) detector for purity assessment and LC-MSD mass spectral detector for identity confirmation at a flow rate of 2 mL/min using a Zorbax SB-C18 (4.6 mm x 30 mm, 2.5 µm) or a Waters Alliance system running at 2 mL/min with an Xterra MS C18 column (4.6 mm x 50 mm, 3.5 µm) with a Model 996 PDA and a Micromass ZQ or Platform II mass spectrometer. Data was captured and processed utilizing the instrument software packages except for MS spectra, which were processed with version 4.0 of MassLynx software. The standard gradient employed on both the Agilent and Waters systems using $H_2O$ and $CH_3CN$ as solvents, each containing 0.1% formic acid, was: 0-0.5 min 5% $CH_3CN$, 0.5-5 min 5->100% $CH_3CN$, 5-7 min 100% $CH_3CN$. UPLC analyses were performed on a Waters Acquity system equipped with PDA and MS detectors at 7 mL/min on an Acquity UPLC BEH C18 column (2.1 mm x 50 mm, 1.7 µm) utilizing a standard gradient also with $H_2O$ and $CH_3CN$, each containing 0.1% formic acid: 0-1 min 5% $CH_3CN$, 1-8 min 5->100% $CH_3CN$, 8-10 min 100% $CH_3CN$. Retention times ($t_R$) relative to that of injection are reported.

Preparative HPLC purifications were performed using the following instrumentation configuration: Waters 2767 Sample Manager, Waters 2545 Binary Gradient Module, Waters 515 HPLC Pumps (2), Waters Flow Splitter, 30-100 mL, 5000:1, Waters 2996 Photodiode Detector, Waters Micromass ZQ, on an Atlantis Prep C18 OBD (19 x 100 mm, 5 µm) or an XTerra MS C18 column (19 x 100 mm, 5 µm) at a flow rate of 30 mL/min using one of the gradients described in the standard method below. The mass spectrometer, HPLC and mass-directed fraction collection were controlled via MassLynx software version 4.0 with FractionLynx.

When necessary, manual flash column chromatography[1] on silica gel (230-400 mesh, MilliporeSigma) and automated medium pressure chromatographic purifications were employed for purification of building block components and, in the latter case, performed on a Biotage Isolera system with disposable silica or C18 cartridges. Solid phase extraction (SPE) was performed utilizing PoraPak™ [Waters, Milford, MA, USA or MilliporeSigma (Supelco), St. Louis, MO, USA], SiliaSep™, SiliaPrep™ and SiliaPrepX™ (SiliCycle, Quebec, QC, Canada) as free media, pre-packed cartridges, or plates as appropriate for the compound(s) being purified.

The expressions "concentrated/evaporated/removed under reduced pressure" and "concentrated/ evaporated/removed *in vacuo*" indicates evaporation of volatile materials utilizing a rotary evaporator under either water aspirator pressure or the stronger vacuum provided by a mechanical oil vacuum pump as appropriate for the solvent being removed or, for multiple samples simultaneously, evaporation of solvents and other volatiles utilizing a centrifugal evaporator system (Genevac HT-24, SP Industries, Warminster, PA, USA).

**Standard Procedure for the Preparation of Alcohol Building Blocks**

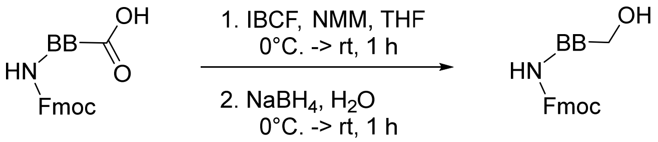

Fmoc-S1 Fmoc-S2

For the transformation of amino acids to the corresponding amino alcohols to be utilized as building block elements for the macrocyclic molecules, a method analogous to those in the literature was employed.[2,3] A solution of the Fmoc-protected amino acid (Fmoc-S1, 1 eq) in THF (6-7 mL/mmol) under nitrogen was cooled to 0°C. in an ice-salt bath, then isobutyl chloroformate (IBCF, 1.05 eq) and 4-methylmorpholine (NMM, 1.05 eq) added dropwise simultaneously or successively over approximately 5 min. The mixture was stirred at 0°C. for 30 min, then at room temperature (rt) for another 30 min. The white precipitate that formed was filtered into a round bottom flask through a pre-washed Celite® pad and rinsed with anhydrous ether. The flask was placed under nitrogen in an ice-bath, and a mixture of sodium borohydride (1.5 eq) in water added in one shot with the neck of the flask left open. Significant gas evolution was observed and the reaction mixture formed a suspension. More water (20 mL) was added, the ice-bath removed, and the reaction stirred rapidly with monitoring by LC-MS and TLC. After 1 h at rt, LC-MS analysis indicated that the reaction was complete. More water was then added and the organic layer extracted with EtOAc (2x). The combined organic layers were washed sequentially with 1 M citric acid (1x), saturated $NaHCO_3$ (1x), water (1x), brine (1x), and dried over anhydrous $MgSO_4$. The mixture was filtered and the filtrate concentrated under reduced pressure to give the alcohol, Fmoc-S2, in 60-80% yield. The product thus obtained was sufficiently pure to be used without further purification for subsequent reactions.

**General Procedure for Oxidation of Alcohol Building Blocks to Aldehydes.**

A number of different oxidation reagents were successfully utilized to convert alcohols, generally Fmoc-protected, to aldehydes for use in attachment by reductive amination. The products were characterized by $^1$H NMR (using the aldehyde C<u>H</u>O as a diagnostic tool) and LC-MS. The following table lists the most common oxidation reagents/methods employed and the types of building blocks on which they were applied:

| ***Oxidation Reagent/Method*** | ***Substrate Types*** |
|---|---|
| $MnO_2$ oxidation[4] | benzylic and pyridine-containing alcohols |
| Pyridine•$SO_3$ (Parikh–Doering oxidation)[5] | alkyl and benzylic alcohols |
| Swern oxidation (DMSO, oxalyl chloride)[6] | benzylic and alkyl alcohols |

| | |
|---|---|
| Dess-Martin periodinane (DMP, 1,1,1-triacetoxy-1,1-dihydro-1,2-benziodoxol-3(1H)-one)[7] | alkyl alcohols |

Additional details on the first two of these methods follow.

**Standard Procedure for the Preparation of Aldehyde Building Blocks Using Pyridine-Sulfur Trioxide Complex**

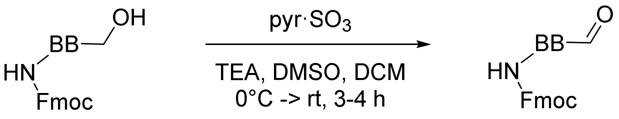

Fmoc-S3 Fmoc-S4

The following procedure was employed for the transformation of Fmoc-protected amino alcohols to the corresponding amino aldehyde building block components for use in the reductive amination attachment procedure. In a round-bottomed flask, the alcohol (1 eq), Fmoc-S3, was dissolved in DCM (4 mL/mmol) and DMSO (1:10 (v/v) with DCM) to leave a clear solution. Triethylamine (TEA, 4 eq) was added and the solution cooled to 0°C under nitrogen. Pyridine sulfur trioxide complex (pyr•$SO_3$, 3 eq) was introduced via syringe as a solution in DMSO (2 mL/mmol pyr•$SO_3$) over 20-30 min and the reaction monitored by TLC and LC-MS until complete, which usually required 3-4 h, while warming to rt. The reaction was cooled back to 0°C in an ice-bath, EtOAc/ether (1:1, 2x reaction volume) added, and the organic layer washed sequentially with cold saturated $NH_4Cl$ (5x), cold saturated $NaHCO_3$ (3x) and brine (2x). More water was added as necessary to dissolve any insoluble material. The aqueous layer was extracted with EtOAc/ether (1:1, 3x). The organic extracts were combined and washed sequentially with 1M $KHSO_4$ (1x), saturated $NH_4Cl$ (2x), water (1x), brine (2x), dried over anhydrous $Na_2SO_4$, filtered and the filtrate concentrated under reduced pressure to give the aldehyde Fmoc-S4. The product thus obtained was usually acceptable for use in subsequent transformations without further purification.

**Standard Procedure for the Preparation of Aldehyde Building Blocks with Manganese Dioxide**

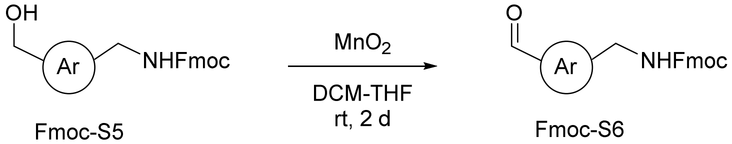

Fmoc-S5 Fmoc-S6

The alcohol, Fmoc-S5 (1 eq) was suspended in DCM (1 mL/mmol) and THF (1 mL/mmol). Manganese dioxide (Strem Chemicals, Newburyport, MA, USA) (1.06 eq) was added and the reaction agitated overnight (o/n) on an orbital shaker at 200 rpm. If the reaction was incomplete, additional quantities of $MnO_2$ were introduced with the mixture agitated for 16 h more after each such addition until starting material was consumed. At that point, the reaction solution was filtered and the residual $MnO_2$ agitated with THF, which was used to wash the material in the filter as well. The $MnO_2$ was washed again with THF and agitated, then passed through the filter. The combined filtrate was dried over anhydrous $MgSO_4$, the solid removed, then the solvent evaporated under reduced pressure to leave the aldehyde, Fmoc-S6, as a solid or syrup. $^1$H-NMR and LC-MS analysis were consistent with the expected product and indicated sufficient purity to proceed with use in macrocycle assembly.

**General Methods for Synthesis of Libraries of Macrocyclic Compounds**

An outline of the general strategy used for the synthesis of the macrocyclic compounds is provided in Supplementary Fig. S6. Although a split-pool strategy is employed for efficiency in the library preparation,[8,9,10] each macrocycle is ultimately obtained as a discrete individual compound through the utilization of multiple, small, meshed, polypropylene containers with snap closures (MiniKans) equipped with a radiofrequency (Rf) tag to permit tracking of the syntheses and identification of the particular structure contained therein prior to resin cleavage.[11]

The general strategy led to macrocyclic compounds containing three, four or five building blocks (compounds **12**, **13**, **14**, respectively in Fig. S6) with additional details of the various steps involved in the assembly provided in the succeeding procedures. The Fmoc/tBu protection strategy[12,13] was employed for their preparation. To initiate the process, the first building block ($BB_1$), as its acid, was attached directly to 2'-chlorotrityl chloride polystyrene resin[14] under basic conditions. The assembly of subsequent components then employed one of three reactions after removal of the Fmoc group: amide bond formation, Mitsunobu-Fukuyama reaction[15] (which required additional manipulations to effect), or reductive amination. Depending on the number of BBs desired for the target macrocycle, the route was terminated after 3, 4 or 5 total components were joined together. At that stage, sequential N-terminal Fmoc removal, cleavage from the resin support, macrocyclization via amide bond formation, and final deprotection of the remaining protecting groups were performed. The crude products thus obtained were roughly purified using SPE procedures followed by preparative HPLC with MS monitoring and collection triggering. For each final macrocycle thus isolated, the HPLC purity (UV) was determined and the identity confirmation by MS. Note that it was found that in certain instances, purification prior to removal of the side chain protection was performed, for example, if separation from side products and reagents was observed in the crude HPLC to be more easily achieved than at the fully deprotected stage.

The QUEST Library of novel macrocyclic compounds used for the initial HTS for this investigation was prepared as described in the patent literature, which also provides the structures of thousands of its individual members.[16,17,18] Briefly, its construction followed analogous steps to those just described: (i) synthesis of the individual multifunctional, appropriately protected, building blocks, including elements for interaction at biological targets and fragments for control and definition of conformation, as well as moieties that can perform both functions; (ii) assembly of the building blocks, typically in a sequential manner with cycles of selective deprotection and attachment, utilizing standard chemical transformations such as amide bond formation, Mitsunobu reaction and its variants, nucleophilic substitution reactions, reductive amination, and metal- and organometallic-catalyzed coupling; (iii) optionally, selective removal of one or more side chain protecting groups was performed, either during the building block assembly or after assembly was completed, then the resin-bound molecule further reacted with one or more additional building blocks to extend the structure at the selectively unprotected functional group(s); (iv) once assembly of the linear intermediate precursor was completed, the N-terminal functional group was deprotected, the linear precursor cleaved from the resin, followed by cyclization of the compounds, which can involve one or more steps, to form the macrocyclic structures; and (v) removal of all remaining protecting groups and purification to provide the desired final macrocycles.

Upon isolation and characterization, the library compounds were stored individually in the form thus obtained (solids, syrups, gums) or dissolved in an appropriate solvent, for example DMSO. In solution, the compounds were distributed into an appropriate array format for ease of use in automated screening assays, such as in microplates or on miniaturized chips. After isolation and to ensure the integrity of the compounds was maintained prior to and in-between any testing, libraries were stored at or below -70°C as 10 mM solutions in 100% DMSO. For use, the plates were allowed to warm to ambient temperature and diluted with buffer, first to a working stock solution, then further to reach the appropriate test concentrations for use in HTS or other assays. It should be noted, however, that no inherent instability of these macrocycles has been observed even upon standing at ambient temperature for extended periods of time.

**General Methods for Solid Phase Chemistry**

For the manipulations described here, polystyrene cross-linked with divinyl benzene (PS-DVB) resins were employed with DMF, DCM and NMP (N-methyl-2-pyrrolidone) as the most common solvents. The volume of the reaction solvent required was generally 3-5 mL per 100 mg resin or approximately 0.04 mL/mg resin. When the term "appropriate amount of solvent" is used in the synthesis methods, it refers to this quantity. Reaction stoichiometry was determined based upon the loading of the starting resin (represents the number of active functional sites, as provided by the supplier, usually as mmol/g). The recommended quantity of solvent roughly amounts to a 0.2 M solution of building blocks at 3-5 eq relative to the initial loading of the resin, except for the initial building block attachment to the resin, for which 2.5 eq was sufficient. Solid phase reactions were conducted in round bottom flasks, solid phase reaction vessels equipped with a fritted filter and stopcock, or Teflon-capped jars. The vessel size was selected so that the solvent/resin mixture only filled ~60% of the container to provide adequate space for the resin to be effectively agitated taking into account these PS-DVB resins swell significantly in organic solvents. Agitations for solid phase chemistry were performed with an orbital shaker (Thermo Scientific, Forma Models 416 or 430, or New Brunswick) at approximately 200 rpm, unless otherwise specified.

The volume of solvent used for the resin washes was a minimum of the same volume as used for the reaction, although more solvent was generally employed to ensure complete removal of excess reagents and other soluble residual by-products (minimally 0.05 mL/mg resin). Each of the resin washes listed in the procedures was performed for a duration of at least 5 min with agitation (unless otherwise specified) in the specific order listed. The number of washings is denoted by "nx" together with the solvent or solution, where n is an integer. In the case of mixed solvent washing systems, they are listed together and denoted solvent 1/solvent 2. After washing, the expression "dried in the usual manner" and analogous expressions mean that the resin was dried first in a stream of nitrogen or argon for 20 min to 1 h, using argon if there was concern over oxidation of the substrate on the resin, and subsequently under vacuum (oil pump usually) until full dryness is attained [minimum 2 h to overnight (o/n)].

**General Procedure for Loading of First Building Block to Resin**

The following procedure was followed for adding the first protected building block to 2'-chlorotrityl chloride resin (75-80 mg, 1.11 mmol/g) was loaded into each MiniKan. The resin was pre-washed with DCM (2x), then dried under a nitrogen flow for 2 h. In a suitable reaction vessel, Fmoc-$NR_1$-$BB_1$-$CO_2H$ (2.5 eq) was dissolved in DCM (0.04 mL/mg resin) and diisopropylethylamine (DIPEA, 5 eq) added, then the solution agitated briefly and the resin introduced. The reaction was agitated o/n at which point the solvent was removed, washed with DMF (2x), then any remaining reactive sites on the resin capped using two successive treatments with MeOH/DIPEA/DCM (2:1:17). After these treatments and removal of the solvent, the resin was washed sequentially with DCM (1x), iPrOH (1x), DCM (1x), ether (1x), then dried in the usual manner.

**Standard Procedure for Monitoring Reaction Progress on the Solid Phase (In-Process Quality Control (QC))**

Since direct methods usually employed for monitoring reaction progress (TLC, GC, HPLC) are not applicable for solid phase reactions, it was necessary to perform cleavage of a minimal amount of material from the resin support in order to determine the progress of each transformation, such as described in the following procedure. Representative duplicate samples in MiniKans specifically for this purpose were included within the library so as not to adversely affect the preparation of any of the target molecules.

For in-process QC, a small amount of resin (a few beads were usually sufficient) was removed from the reaction vessel, then washed successively with DMF (2x), iPrOH (1x), DCM (2x), ether (1x), dried briefly with nitrogen flow, then treated with 250 µL 20% hexafluoroisopropanol (HFIP)/DCM for 10-20 min, and concentrated with a stream of nitrogen. To the crude residue obtained was added 1 mL MeOH (if necessary to solubilize the fully protected crude intermediate compounds, a small amount of DMSO or THF was also employed), the solution filtered through a 45 µm HPLC filter, or, alternatively, a plug of cotton in a small pipette, and the filtrate analyzed by HPLC or HPLC-MS. For solid phase reactions involving amines, the Kaiser (ninhydrin) test was also used to monitor progress.[19]

**General Procedure for Fmoc Deprotection**

In an appropriate vessel, a solution of 20% piperidine (pip) in DMF was prepared, an excess added to the resin and the mixture agitated for 30 min. The reaction solution was removed, then this treatment repeated. After draining of the solvent, the resin was washed sequentially with: DMF (2x), iPrOH (1x), DMF (1x), iPrOH (1x), THF (1x), DCM (1x), ether (1x), then dried in the standard manner.

In order to minimize the potential of diketopiperazine formation when N-alkylated-amino acids were present in the $BB_1$ position, 50% pip/DMF was used for Fmoc-deprotection of $BB_2$ and the procedure further modified as follows: agitation of the deprotection solution with the resin was performed for only 5-7 min, then the solvent drained. DMF was added to rinse the resin, the mixture agitated quickly and the solvent removed. The washing sequence as described above was then executed.

**General Procedure for Attachment of Amines to Acids**

To an appropriate vessel, DIPEA (7 eq) in NMP was added to the acid-containing building block (3.5 eq), HATU (1-[bis(dimethylamino)methylene]-1H-1,2,3-triazolo[4,5-b]pyridinium 3-oxide hexafluorophosphate, 3.5 eq) in NMP (0.04 mL/mg resin). The mixture was agitated vigorously until all the reactants were solubilized and then added to the resin containing the free amine component. After o/n agitation, the solution was removed and the resin washed sequentially with: DMF (2x), iPrOH (1x), DMF (1x), iPrOH (1x), THF (1x), DCM (1x), ether (1x), then dried in the usual way. For attachment of the third building block (Fmoc-$NR_3$-$BB_3$-$CO_2H$) and beyond, 5 eq of the acid-containing building block and 5 eq of the coupling agent with 10 eq of DIPEA were employed. For any attachment of this type, if the acid building block is one known to require repeated treatment for optimal results (i.e. N-alkylated and other sterically hindered amino acids), half of the indicated equivalents was used for each of two treatments.

**Specialized Procedure for Attachment of Amines to Certain Acids**

When an acid-building block known to be prone to racemization (i.e. Fmoc-phenylglycine and derivatives) was added, modified conditions were employed. In such a case, the acid-containing building block (3.5 eq) and COMU [(1-cyano-2-ethoxy-2-oxoethylidenaminooxy)-dimethylamino-morpholino-carbenium hexafluorophosphate, 3.5 eq] in NMP were combined and 2,6-lutidine (DMPy, 2,6-dimethylpyridine, 5.6 eq) added. After a brief agitation (≤ 1 min), the resin with the free amine component was introduced into the mixture, which was then agitated for 5.5 h. The solution was removed and the resin washed sequentially with: DMF (2x), iPrOH (1x), DMF (1x), iPrOH (1x), DMF (1x), iPrOH (1x), THF (1x), DCM (2x), ether (1x), then dried in the standard manner.

**General Procedure for Attachment of Building Blocks by Reductive Amination using Borane**

The N-protected aldehyde (1.0-2.0 eq) was dissolved in MeOH/DCM/TMOF (trimethyl orthoformate) (2:1:1) and the resulting solution added to the amine-containing resin, then agitated for 0.5 h. To this was introduced borane-pyridine complex (BAP, 4 eq) and the reaction mixture agitated for 15 min, then carefully vented to release built-up pressure. The agitation was continued for another 45 min, vented again, then allowed to proceed o/n. If in-process QC indicated the reaction was not complete, more BAP (2 eq) was added and the mixture agitated again o/n. Once complete, the solvent was removed and the resin washed sequentially with DMF (2x), iPrOH (1x), THF/MeOH (3:1, 1x), DCM/MeOH (3:1, 1x), iPrOH (1x), DMF (1x), DCM (2x), ether (1x), then dried in the usual way.

**General Procedure for Attachment of Building Blocks by Reductive Amination using Sodium Triacetoxyborohydride**

As the preferred method for attachment of aromatic aldehydes, sodium triacetoxyborohydride was employed in the reductive amination process as follows: 1.2-1.5 eq of the Fmoc-protected aldehyde was dissolved in DCM, the amine-containing resin added, then the mixture agitated for 1.5-3.0 h. To this was introduced $NaBH(OAc)_3$ (5 eq) and the reaction agitated o/n with venting at 0.5 h and 1 h. If in-process QC indicated that free starting amine remained on the resin, additional aldehyde (0.6-0.7 eq) was included as part of the repeat treatment along with the reducing agent. Once the reaction was completed, the solvent was removed, then the resin washed sequentially with DMF (2x), iPrOH (1x), MeOH (3:1, 1x), DCM/MeOH (3:1, 1x), iPrOH (1x), THF (1x), DCM (1x), ether (1x) and dried in the standard manner.

**General Procedure for Attachment of Building Blocks by Reductive Amination using Sequential Sodium Cyanoborohydride and BAP Treatment**

For certain benzylic aldehydes, a sequential Borch and BAP reduction process was found to be beneficial. In the first step, the Fmoc-protected aldehyde (3 eq) in NMP/TMOF (1:1) containing 0.5% glacial acetic acid was added to the resin in an appropriate reaction vessel and agitated for 30 min. To the mixture, $NaBH_3CN$ (10 eq) was introduced and the reaction shaken for 10 min, then pressure released and agitation continued o/n. Once in-process QC showed the transformation was complete, solvent was removed and the resin washed sequentially with: DMF (2x), iPrOH (1x), DMF (1x), iPrOH (1x), DCM (2x), ether (1x). If an incomplete reaction was indicated, the solvent was removed and the resin suspended in MeOH/DCM/TMOF (2:1:1). To this was added BAP (2-3 eq) and agitation allowed to proceed for 4 h. The solvent was removed and the resin washed sequentially with: DMF (2x), THF (1x), iPrOH (1x), DCM (1x), THF/MeOH (3:1, 1x), DCM/MeOH (3:1, 1x), DCM (2x), ether (1x), then dried in the usual manner.

**Standard Procedure for Building Block Attachment using Fukuyama-Mitsunobu Reaction**

The resin bound activated amine component was obtained either by attachment of the N-nosylated building block or prepared from an N-Fmoc resin component via N-deprotection using the standard deprotection method and then installation of the N-nosyl moiety as illustrated in Supplementary Fig. S7. This latter transformation was accomplished as follows: 2-nitrobenzenesulfonyl chloride (Nos-Cl, 4 eq) was dissolved in THF:NMP (2:1) and 2,4,6-collidine (10 eq) added, then the resin containing the deprotected amine moiety introduced and the mixture agitated for 1-2 h. The solution was removed and the resin washed sequentially with: DMF(2x), iPrOH (1x), DMF (1x), iPrOH (1x), THF (1x), DCM (1x), ether (1x), DCM (1x), ether (1x), then dried under vacuum.

With the Nos group in place, the following method was used to alkylate the nitrogen under Fukuyama-Mitsunobu conditions[15] for connection of a hydroxy-containing building block (alcohol or phenol, R-OH) to the activated amine component on resin. This procedure was utilized for preparing N-methyl and other N-alkyl components for which the respective individual building block was not commercially available or otherwise difficult to access. The building block (5 eq) was dissolved in THF, 3 Å molecular sieves (200 mg/MiniKan) added and the mixture agitated for 20-30 min before the activated amine containing resin was introduced. After 1.5 h, the solution was cooled to 0°C and the $PPh_3$-DIAD adduct (5 eq, prepared as described below) added. The reaction was agitated o/n while being allowed to warm to rt. The resin was removed by filtration and washed sequentially with: DMF (1x), i-PrOH (1x), DMF (1x), i-PrOH (1x), THF (1x), DCM (1x), ether (1x) then dried in the usual manner. When methanol was used as the alcohol component to prepare N-methylated derivatives, the molecular sieves were not included and the resin added 5 min after the $PPh_3$-DIAD adduct.

After alkylation, the nosyl group was typically removed using the standard method below, then the next building block added or, if the building block assembly was concluded, the precursor was cleaved from the resin and subjected to macrocyclization. Alternatively, in selected cases, the N-Nos group was maintained and its cleavage delayed until the end of the building block assembly or even until after the macrocyclization, since it provided protection of the backbone amide and served to prevent side reactions at that site.[20]

**Standard Procedure for Nosyl Deprotection**

The N-Nos moiety was removed if further chemistry was performed on that nitrogen atom. To effect this, a solution of 2-mercaptoethanol (10 eq), DBU (1,8-diaza-bicyclo[5.4.0.]undec-7-ene, 5 eq) in DMF was prepared, then half added to the resin and the mixture agitated for 8-15 min. The resin was filtered and washed with DMF (3x), then the treatment repeated. The resin was again filtered and washed sequentially with: DMF (2x), iPrOH (1x), DMF (1x), iPrOH (1x), THF (1x), DCM (2x), ether (1x), then dried in the usual way.

**Standard Procedure for the Synthesis of $PPh_3$-DIAD Adduct**

This reagent was prepared essentially as previously reported.[21] In a round bottom flask under nitrogen, diisopropyl azodicarboxylate (DIAD, 1 eq) was added dropwise to a solution of triphenylphosphine ($PPh_3$, 1 eq) in THF (0.4 M) at 0°C, then the reaction stirred for 30 min at that temperature. The resulting precipitate was collected on a cooled, medium porosity glass-fritted filter, the solid washed with cold THF (DriSolv grade) to remove any color, then with anhydrous ether. The white powder was dried *in vacuo* (vacuum pump) and stored under nitrogen in the freezer. It was removed from cold storage shortly before each intended use.

**General Procedure for Cleavage from 2'-Chlorotrityl Resin**

Resin was liberated from the individual MiniKans and transferred into bottom-filtered reaction tubes compatible with the 48 well format of the Mettler Toledo/Bohdan Miniblocks (now available from SiliCycle). To each tube was added 20% HFIP/DCM (1.5 mL), the block covered and agitated on an orbital shaker (460-500 rpm) for 1-2 h. The solution was drained into 7.5 mL tubes arrayed to collect from each of the 48 wells in the Miniblock (pushed out with nitrogen) and the process repeated with an additional quantity of 20% HFIP/DCM (1.5 mL/tube). The resin was then rinsed with DCM (1 mL, again pushed out with nitrogen) and the solvents in the receiving tubes evaporated *in vacuo* (Genevac). The crude material was obtained as solids, semi-solids, syrups or gums.

**General Procedures for Macrocyclization**

*Different procedures for cyclization and subsequent SPE processing (using pre-packed or manually packed cartridges and microplates) were employed depending on the nature of the backbone of the macrocycle product as either basic (containing a secondary or tertiary amine) or neutral (lacking such an amine moiety).*

For macrocycles possessing basic backbones: A solution of 3-(diethoxyphosphoryloxy)-1,2,3-benzotriazin-4(3H)-one (DEPBT, 1.2-1.6 eq) and DIPEA (2.0-2.4 eq) in 25% NMP/THF (0.03 mL/mg original resin) was prepared and added to the crude residue from the prior resin cleavage step. The mixture was then agitated for 24 h at rt. In cases where the compounds were poorly soluble, the residue was dissolved first in NMP, then the DEPBT and DIPEA in THF added to the solution. After the o/n agitation, each reaction mixture was loaded using THF onto PoraPak (p-toluenesulfonic acid, Waters or MilliporeSigma) in cartridges or 48-well 7.5 mL filter plate tubes preconditioned with DCM. Each cartridge/well was sequentially washed with DMF (2x), THF (2x), DCM (2x), then the desired cyclized material eluted with 3.5 M $NH_3$/MeOH (1x), followed by (3.5 M $NH_3$/MeOH)/DCM (1:1, 1x). The eluent was removed under reduced pressure (Genevac). LC-MS of the resulting residues confirmed the presence of the expected products and the absence of the linear starting materials.

The residue for each sample was then individually passed through Si-Carbonate (40-63 µm, 60 Å, SiliCycle) functionalized gel in 48-well filter plates (~270 mg per well), preconditioned with DCM (1 x 1 mL/well) followed by 12% MeOH/DCM (1x), then loaded in 10% MeOH/DCM with washes utilizing the same solvent, followed by elution with 10% MeOH/DCM (3x). Eluents were evaporated *in vacuo* (Genevac).

For macrocycles possessing neutral backbones: DEPBT (1.6 eq) was dissolved in DMF (anhydrous, amine free) to produce a colorless solution which was added to a separate vial containing each crude sample from the resin cleavage. To the resulting homogeneous yellow solutions was added DIPEA (1.9 eq) with the mixture becoming slightly darker yellow over time, but remained homogeneous. The DMF was slowly evaporated with nitrogen flow (2-3 d, rt), then the residue dissolved in 10% MeOH/DCM. These solutions were loaded onto Si-Carbonate in cartridges or 7.5 mL 48-well plates with filter bottom (1.5 g/well), each cartridge/well pre-conditioned with DCM, then 12% MeOH/DCM. The solutions (or suspensions) from each vial were transferred with 12% MeOH/DCM, then eluted with this solvent (3x). After evaporation of the eluents under reduced pressure, the residue was subjected to a second Si-Carbonate purification conducted in an equivalent manner to the first except that the final elution was done with a larger volume of 12% MeOH/DCM. LC-MS of each sample confirmed the presence of the expected cyclized product with none of the linear precursor observed. The amount of dimer was typically greater in these neutral backbone compounds requiring additional care during subsequent prep-LC purification.

Depending on the separation of the protected monomer and protected dimer seen in the LC-MS analyses of the crude cyclized products, certain prep-LC purifications were conducted prior to final deprotection, although the majority were performed on fully deprotected macrocycles.

**Standard Procedures for Final Protecting Group Removal**

The method of deprotection depended on the nature of the protecting groups present on the side chains of the individual macrocycle using the following guidelines:

a) Macrocycles containing Arg, Ser, Thr, Tyr, Gln, Asn, Asp, Glu: 91% TFA, 4% DCM, 2.5% triisopropylsilane (TIPS), 2.5% $H_2O$

b) Macrocycles containing only Lys, Orn, Trp: 50% TFA, 47% DCM, 3% TIPS

c) Macrocycles containing a double bond: 50% TFA, 45%DCM, 5% $H_2O$ (to avoid reduction of the alkene)

The deprotection cocktails indicated (2 mL/cpd) were added to the crude cyclized products obtained after the SPE processing and the resulting mixtures agitated for ~2 hr. Once HPLC indicated deprotection had been completed, the volatiles were removed *in vacuo* (Genevac), 95% DMSO/water (1.5 mL) added to each of the residues and then agitated for 5 h at rt. Analyses were performed on the crude deprotected materials prior to being subjected to preparative HPLC purification using the standard methods below.

**Standard Methods for Preparative HPLC Purification**

Prep-LC purification was performed on the solutions obtained from the deprotection reactions using the instrumentation equipped with mass-triggered fraction collection outlined earlier. The preparative gradients utilized, with aqueous buffer (10 mM ammonium formate, pH 4) and MeOH as eluting solvents were:

P1: 0-2 min 11% MeOH, 2-8 min 11->98% MeOH, 8-9.7 min 98% MeOH, 9.7-10 min 98->50% MeOH (alternative gradients starting at 20, 30 or 40% MeOH were also employed depending on the results of the HPLC analysis)

P5: 0-2 min 11% MeOH, 2-12 min 11->98% MeOH, 12-14.7 min 98% MeOH, 14.7-15 min 98->30% MeOH (alternatives starting at 20% MeOH or of 10 or 12 min total length instead of 15 min or with an initial step of 0-3 min and 12 min total length were also utilized)

Method P5 and its alternatives were employed when a sample required additional purification after the initial preparative run. Lower flow rates (i.e. 20-25 mL/min) were occasionally utilized with concomitant lengthening of the gradient run time.

Fractions indicated by MS analysis to contain the desired pure product were evaporated under reduced pressure (Genevac) or, alternatively, lyophilized. Compounds thus obtained were then analyzed by HPLC/UPLC-MS-

UV analysis for purity assessment and identity confirmation. The use of ammonium formate buffer resulted in the macrocyclic compounds being obtained as their formate salt forms.

**Characterization of Macrocyclic Compounds**

The following macrocycles are representative of the chemotypes identified as the initial hits from the original HTS of the QUEST Library, as well as the validated hits from the subsequent focused library. These selections include the most active PPI stabilizers found from these investigations.

**CY000970**

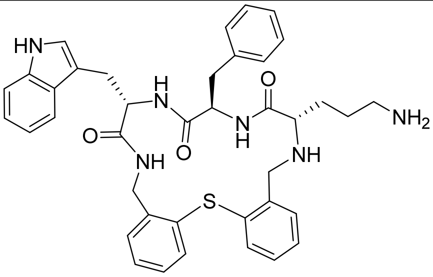

CY000970

$^1$H NMR (DMSO-$d_6$, 500 MHz): δ 1.23-1.45 (br m, 4H), 2.53-2.68 (m, 4H), 2.90 (dd, $J$ = 10.0; 15.0 Hz, 1.5H), 3.07-3.13 (m, 2.5H), 3.40 (d, $J$ = 15.0 Hz,2H), 3.81 (d, $J$ = 15.0 Hz, 2H estimated since a very broad water peak centered at 4.20 ppm complicated accurate integration of other resonances in the 3.7-4.7 range), 4.16 (dd, $J$ = 5.0; 10.0 Hz, 1H estimated), 4.34 (dd, $J$ = 5.0; 15.0 Hz, 2H estimated), 4.48 (dt, $J$ = 5.0; 10.0 Hz, 1H estimated), 4.59 (br q, $J$ = 5.0 Hz, 1H estimated), 6.78-6.82 (m, 1H), 6.95-7.39 (m, 14H), 7.54 (br d, $J$ = 5.0 Hz, 1H), 7.61 (br d, $J$ = 5.0 Hz, 1H), 7.83 (br t, $J$ = 5.0 Hz, 1H), 8.38-8.53 [br s, total 5H, overlapped with 8.47 (dd, $J$ = 5.0; 20.0 Hz)], 10.86 (s, 1H)

LC-MS: $t_R$ 2.47 min, 100% (UV, 220 nm), [$(M+H)^+$] 675.30

**CY001110**

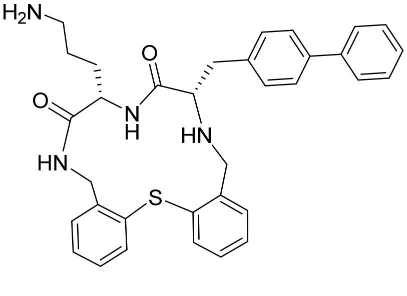

CY001110

$^1$H NMR (DMSO-$d_6$, 500 MHz): δ 1.40-1.50 (m, 2H), 1.54-1.63 (m, 1H), 1.71-1.80 (m, 1H), 2.72 (br t, $J$ = 7.5 Hz, 2H), 2.90 (dq, $J$ = 5.0; 15.0 Hz, 2H), 3.23 (t, $J$ = 7.5 Hz, 1H), 3.38 (d, $J$ = 10.0 Hz, 2H), 3.59 (d, $J$ = 10.0 Hz, 2H), 4.08 (dd, $J$ = 2.5; 15.0 Hz, 1H), 4.26 (q, $J$ = 10.0 Hz, 1H), 4.70 (dd, $J$ = 5.0; 15.0 Hz, 1H), 6.31 (dd, $J$ = 2.5; 5.0 Hz, 1H), 7.04-7.11 (symmetric m, 2H), 7.24-7.67 (m, 15H), 8.18 (d, $J$ = 10.0 Hz, 1H), 8.27 (br dd, $J$ = 5.0; 10.0 Hz, 1H), 8.42 (br s, 1H)

LC-MS: $t_R$ 2.26 min, 100% (UV, 220 nm), [$(M+H)^+$] 565.20

**CY001896**

$^{1}$H NMR (DMSO-d$_6$, 500 MHz): δ 1.21-1.57 (m, 10H), 2.57-2.66 (m, 2H), 2.72 (br t, *J* = 5.0 Hz, 2H), 2.82 (dd, *J* = 7.5; 15.0 Hz, 1H), 2.87 (t, *J* = 7.5 Hz, 1H), 2.98 (dd, *J* = 5.0; 15.0 Hz, 1H), 3.71 (dd, *J* = 15.0; 65.0 Hz, 2H), 4.01-4.12 (m, 2H), 4.25 (br q, *J* = 5.0 Hz, 1H), 4.44 (dd, *J* = 5.0; 15.0 Hz, 1H), 6.63 (d, *J* = 10.0 Hz, 2H), 6.77-6.80 (m, 1H), 6.92 (d, *J* = 10.0 Hz, 2H), 7.12 (dd, *J* = 2.5; 10.0 Hz, 1H), 7.19-7.27 (m, 3H), 7.37 (dt, *J* = 2.5; 7.5 Hz, 1H), 7.40-7.43 (m, 1H), 7.61 (br d, *J* = 10.0 Hz, 1H), 7.99 (dd, *J* = 5.0; 10.0 Hz, 1H), 8.32 (br d, *J* = 5.0 Hz, 1H), 8.43 (br s, 1H) overlapping with 8.46 (br d, *J* = 10.0 Hz, 1H)

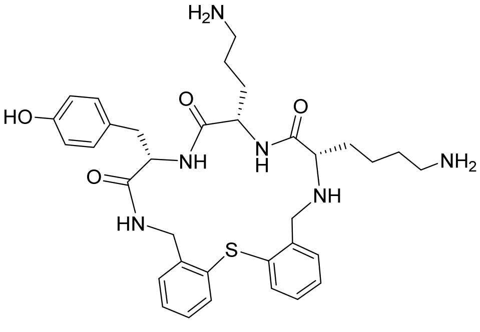

CY001896

UPLC: $t_R$ 2.73 min, 99.1% (UV, 220 nm), 0.9% diastereomer ($t_R$ 2.84 min)

LC-MS: $t_R$ 1.95 min, 100% (UV, 220 nm), [$(M+H)^+$] 633.30

**CY001951**

$^{1}$H NMR (DMSO-d$_6$, 500 MHz): δ 1.13-1.59 (m, 11H), 2.71 (br t, *J* = 7.5 Hz, 2H), 2.82 (dd, *J* = 10.0; 15.0 Hz, 1H), 2.88 (t, *J*= 7.5 Hz, 1H), 2.90-3.04 (m, 4H), 3.63 (d, *J* = 15.0 Hz, 2H), 3.79 (d, *J* = 15.0 Hz, 2H), 4.02 (dd, *J* = 2.5; 15.0 Hz, 1H) partial overlap with 4.08 (br q, *J* = 7.5 Hz, 1H), 4.25 (br q, *J* = 7.5 Hz, 1H), 4.46 (dd, *J* = 5.0; 10.0 Hz, 1H), 6.64 (d, *J* = 10.0 Hz, 2H), 6.78-6.82 (m, 1H), 6.92 (d, *J* = 10.0 Hz, 2H), 7.12 (d, 1*J* = 10.0 Hz, 1H), 7.19-7.27 (m, 3H), 7.37 (t, *J* = 10.0 Hz, 1H), 7.40-7.43 (m, 1H), 7.61 (d, *J* = 10.0 Hz, 1H), 7.90 (br s, not integrated), 7.96 (dd, *J* = 5.0; 5.0 Hz, 1H), 8.20 (d, *J* = 5.0 Hz, 1H), 8.42 (s) overlapping with 8.45 (d, *J* = 10.0 Hz, total 3H)

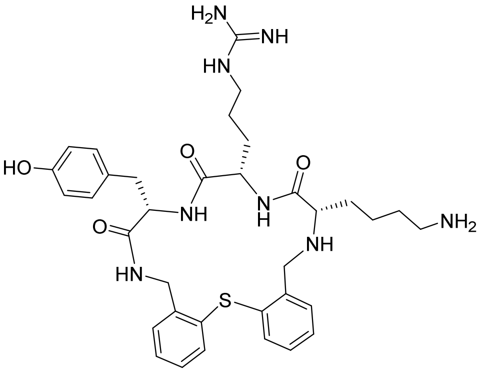

CY001951

LC-MS: $t_R$ 2.02 min, 100% (UV, 220 nm), [$(M+H)^+$] 675.30

**CY002028**

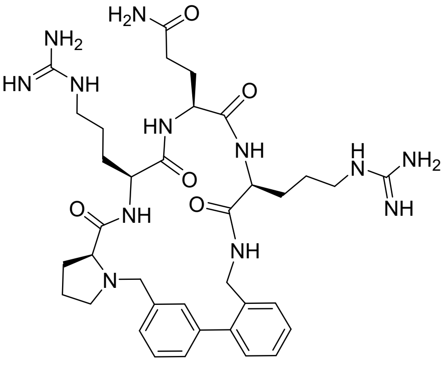

CY002028

$^{1}$H NMR (DMSO-$d_6$, 500 MHz): δ 1.22-1.93 (m, 14H), 2.05-2.18 (m, 3H), 2.28 (br q, *J* = 5.0 Hz, 1H), 2.80-2.87 (m, 3H), 3.01-3.10 (m, 4H), 3.34 (d, *J* = 10.0 Hz, 2H), 3.72-3.77 (m, 2H estimated due to very broad water peak centered at ~3.70 ppm complicating accurate integration of other resonances in 3.3-4.3 range), 3.95-4.01 (m, 2H estimated), 4.05 (d, *J* = 15.0 Hz, 2H estimated), 4.40-4.46 (m, 2H), 6.80 (br s, 1 H), 7.14 (dd, *J* = 2.5; 10.0 Hz, 1H), 7.20 (d, *J* = 5.0 Hz, 1H), 7.26-7.42 (m, 8H), 7.50-7.70 (br m, 4H) overlapping 7.63 (s), 8.02 (v br d, *J* = 10.0 Hz, 1H), 8.35 (br s, 3H), 8.47 (br d, 1H, *J* = 5.0 Hz), 8.67 (br s, 1H), 8.87 (br s, 1H)

LC-MS: $t_R$ 1.97 min, 100% (UV, 220 nm), [$(M+H)^+$] 733.50

**CY007183**

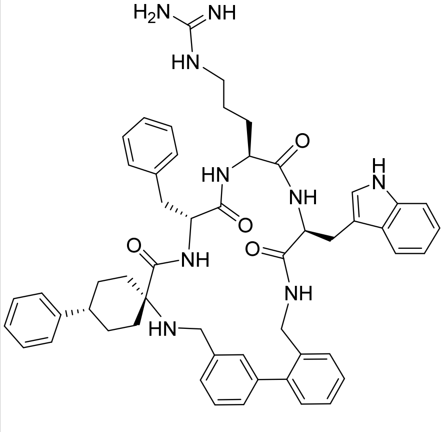

CY007183

$^{1}$H NMR (DMSO-$d_6$, 500 MHz): δ 1.06-1.14 (m, 1H), 1.16-1.27 (m, 1H), 1.32-1.40 (m, 1H), 1.52-1.65 (m, 3H), 1.66-1.92 (m, 6H), 2.86-2.95 (m, 3H), 2.98-3.05 (m, 2H), 3.24 (dd, *J* = 5.0; 15.0 Hz, 1H), 3.45 (unsymmetrical dd, *J* = 15.0; 50.0 Hz, 4H estimated due to overlap with broad water peak), 3.72-3.77 (m, 1H), 4.16 (unsymmetrical dd, *J* = 5.0; 15.0 Hz, 1H), 4.25 (unsymmetrical dd, *J* = 5.0; 15.0 Hz, 1H), 4.40-4.47 (m, 1 H), 4.65 (q, *J* = 5.0 Hz, 1H), 6.95 [dt, *J* = 1.0 (estimated); 5.0 Hz, 1H], 7.01-7.07 (m, 4H), 7.10-7.42 (m, 17 H), 7.49 (d, *J* = 5.0 Hz, 1H), 7.57 (br s, not integrated), 7.75 (d, *J* = 10.0 Hz, 1H), 7.92 (t, *J* = 5.0 Hz, 1H), 8.15 (d, *J* = 5.0 Hz, 1H), 8.42 (br s, 1H), 8.61 (br t, J = 5.0 Hz, 1H), 8.72 (d, *J* = 5.0 Hz, 1H), 10.85 [barely resolved d, *J* = 1.0 Hz (est.), 1H]

LC-MS: $t_R$ 2.98 min, 100% (UV, 220 nm), [$(M+H)^+$] 886.60

**CY007274**

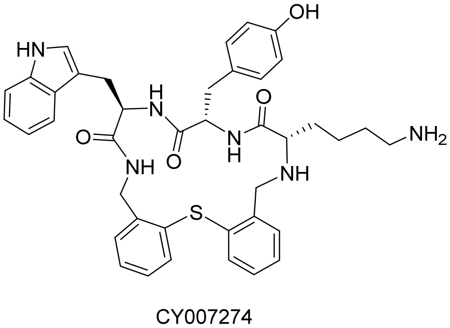

CY007274

$^1H$ NMR (DMSO-$d_6$, 500 MHz): δ 0.76-0.84 (m, 1H), 0.96-1.05 (m, 1H), 1.20-1.30 (m, 1H), 1.31-1.40 (m, 3H), 2.36-2.48 (m, 2H, partial overlap with DMSO-$d_5$ peaks), 2.56-2.64 (m, 3H), 2.94 (dd, *J* = 10.0; 15.0 Hz, 2H), 3.13 (dd, *J* = 5.0; 20.0 Hz, 2H), 3.64 (d, *J* = 15.0 Hz, 2H), 3.91 (d, *J* = 15.0 Hz, 2H), 4.13 (dd, *J* = 5.0; 15.0 Hz, 2H), 4.44 (dd, *J* = 5.0; 15.0 Hz, 1H), 4.50 (dt, *J* = 5.0; 10.0 Hz, 1H), 4.59 (dt, *J* = 5.0; 10.0 Hz, 1H), 6.40-6.43 (m, 1H), 6.56 (d, *J* = 10.0 Hz, 1H), 6.77 (d, *J* = 10.0 Hz, 2H), 6.98 [barely resolved dt, *J* = 1.0 (est.); 7.5 Hz, 1H], 7.05 [barely resolved dt, *J* = 1.0 (est.); 7.5 Hz, 1H], 7.09-7.15 (m, 3H), 7.25-7.28 (m, 2H), 7.30 (d, *J* = 10.0 Hz, 1H), 7.33-7.37 (m, 1H), 7.41-7.46 (m, 1H), 7.57 (d, *J* = 10.0 Hz, 1H), 7.63 (d, *J* = 10.0 Hz, 1H), 7.70 (d, *J* = 10.0 Hz, 1H), 7.90 (br t, *J* = 5.0 Hz, 1H), 8.40 (br s, 2H), 8.53 (d, *J* = 10.0 Hz, 1H), 10.85 [barely resolved d, *J* = 1.0 Hz (est.), 1H],

LC-MS: $t_R$ 2.37 min, 96.6% (UV, 220 nm), $[(M+H)^+]$ 705.30; impurity $t_R$ 2.51 min, 3.4% (UV, 220 nm), $[(M+18+H)^+]$ 723.30

**CY007288**

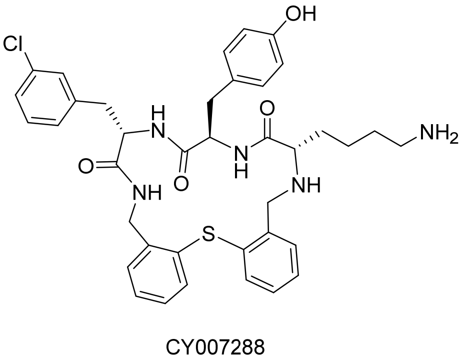

CY007288

$^1H$ NMR (DMSO-$d_6$, 700 MHz): δ 0.73-0.80 (m, 1H), 0.98-1.05 (m, 1H), 1.17-1.24 (m, 1H), 1.25-1.31 (m, 1H), 1.32-1.43 (m, 2H), 2.33 (dd, *J* = 7.0; 14.0 Hz, 1H), 2.44 (dd, *J* = 7.0; 14.0 Hz, 1H), 2.59 (br t, *J* = 7.0 Hz, 2H), 2.78 (dd, *J* = 7.0; 14.0 Hz, 1H), 3.05 (dd, *J* = 7.0; 14.0 Hz, 1H), 3.08 (t, *J* = 7.0 Hz, 1H), 3.37 (d, *J* = 14.0 Hz, 1H), 3.79 (d, *J* = 14.0 Hz, 1H), 4.13 (dd, *J* = 7.0; 14.0 Hz, 1H), 4.39 (dd, *J* = 7.0; 14.0 Hz, 1H), 4.46-4.52 (m, 2H), 6.63 (d, *J* = 7.0 Hz, 2H), 6.91 (partially resolved dd, *J* = 7.0; 7.0 Hz) overlapped with 6.94 (d, *J* = 7.0 Hz, total 3H), 7.00 (d, *J* = 7.0 Hz, 1H), 7.17-7.31 (m, 8H), 7.37 (br s, 1H), 7.43-7.46 (m, 1H), 7.57 (d, *J* = 7.0 Hz, 1H), 7.85 (br t, *J* = 7.0 Hz, 1H), 8.35-8,.47 [(br s), total 3H, overlapping 8.42 (d, *J* = 7.0 Hz)], 8.67 (d, *J* = 7.0 Hz, 1H)

$^{13}C$ NMR (DMSO-$d_6$, 176 MHz): δ 22.4, 27.3, 32.7, 36.2, 36.4, 38.7, 42.2, 48.5, 53.4, 54.2, 61.1, 114.9, 126.3, 127.0, 127.48, 127.52, 127.7, 128.1, 128.2, 128.9, 129.2, 129.8, 130.0, 130.6, 130.7, 131.7, 132.2, 132.6, 135.1, 137.6, 140.8, 141.0, 156.2, 165.4 (br), 170.1, 172.0, 174.3

LC-MS: $t_R$ 2.50 min, 100% (UV, 220 nm), $[(M+H)^+]$ 700.30

HRMS (ESI-TOF, $[M+H]^+$, *m/z*): Calculated for $C_{38}H_{43}ClN_5O_4S$ 700.2719, Found 700.2731

**CY007345**

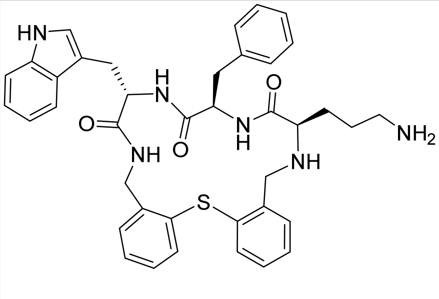

CY007345

$^{1}$H NMR (DMSO-d$_6$, 500 MHz): δ 1.21-1.47 (m, 4H), 2.53-2.66 (m, 3H), 2.69 (br t, *J* = 5.0 Hz, 1H), 2.90 (dd, *J* = 10.0; 15.0 Hz, 1H), 3.08 (dd, *J* = 5.0; 15.0 Hz, 1H), 3.78 (d, *J* = 15.0 Hz, 2H), 3.87 (d, *J* = 15.0 Hz, 2H), 4.14 (dd, *J* = 5.0; 10.0 Hz, 1H), 4.41 (dd, *J* = 5.0; 15.0 Hz, 1H), 4.49 (dt, *J* = 5.0; 10.0 Hz, 1H), 4.64 (br q, *J* = 5.0 Hz, 1H), 6.43-6.49 (m, 1 H), 6.90-6.94 (m, 2H), 6.96-7.01 (m, 1H), 7.03-7.08 (m, 2H), 7.08-7.16 (m, 5H), 7.23-7.37 (m, 4H), 7.40-7.45 (m, 1H), 7.53 (d, *J* = 10.0 Hz, 1H), 7.62 (d, *J* = 10.0 Hz, 1H), 7.85 (d, *J* = 5.0 Hz, 1H), 7.91 (approximates t, *J* = 5.0 Hz, 1H), 8.42 (br s, 1H) partially overlapping with 8.49 (d, *J* = 5.0 Hz, 1H), 10.84 (s, 1 H)

LC-MS: $t_R$ 2.58 min, 100% (UV, 220 nm), [(M+H)$^+$] 675.30

**CY007424**

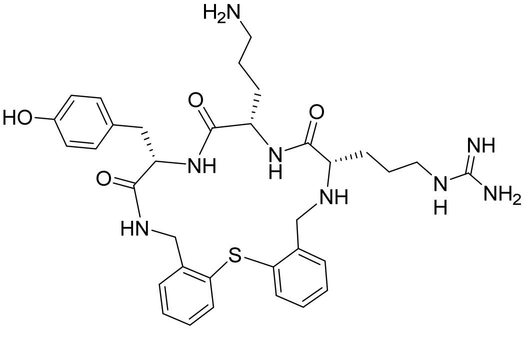

CY007424

$^{1}$H NMR (DMSO-d$_6$, 700 MHz): δ 1.22-1.31 (m, 1H), 1.39-1.59 (m, 7H), 2.60-2.70 (m, 2H), 2.82 (dd, *J* = 7.0; 14.0 Hz, 1H), 2,89 (t, *J* = 7.0 Hz, 1H), 2.98 (dd, *J* = 7.0; 14.0 Hz, 1H) partially overlapped with 2.97-3.08 (m, 4H), 3.65 (d, *J* = 14.0 Hz, cannot obtain accurate integration due to large water peak, 2H estimated), 3.78 (d, *J* = 14.0 Hz, cannot obtain accurate integration due to large water peak, 2H estimated), 4.04 (dd, *J* = 7.0; 14.0 Hz, 1H), 4.06-4.12 (m, 1H), 4.25 (q, *J* = 7.0 Hz, 1H), 4.44 (dd, *J* = 7.0; 14.0 Hz, 1H), 6.64 (d, *J* = 7.0 Hz, 2H), 6.76-6.79 (m, 1H), 6.93 (d, *J* = 7.0 Hz, 2H), 7.14 (d, *J* = 7.0 Hz, 1H), 7.18-7.23 (m, 2H) partial overlap with 7.25 (t, *J* = 7.0 Hz, 1H), 7.38 (t, *J* = 7.0 Hz, 1H), 7.40-7.43 (m, 1H), 7.61 (d, *J* = 7.0 Hz, 1H), 7.88 (v br s, 2H), 7.99 (partially resolved t, *J* = 7.0 Hz, 1H), 8.31 (d, *J* = 7.0 Hz, 1H), 8.40 (br-s, 3H), 8.50 (d, *J* = 7.0 Hz, 1H), 8.95 (br s, 0.5H)

$^{13}$C NMR (DMSO-d$_6$, 176 MHz): δ 24.2, 25.3, 29.2, 30.6, 35.8, 38.4, 40.4, 42.2, 48.3, 53.7, 55.4, 62.5, 115.0, 126.7, 127.7, 127.8, 128.2, 128.78, 128.81, 129.6, 130.1, 131.8, 131.9, 132.3, 135.4, 137.1, 141.6, 156.0, 157.5, 166.2, 170.2, 171.6, 174.2

LC-MS: $t_R$ 2.00 min, 100% (UV, 220 nm), [(M+H)$^+$] 661.30

HRMS (ESI-TOF, [M+H]$^+$, *m/z*): Calculated for $C_{34}H_{45}N_8O_4S$ 661.32790, Found 661.32917

**CY007455**

$^1$H NMR (DMSO-d$_6$, 500 MHz): δ 1.20-1.29 (m, 2H), 1.43-1.53 (m, 2H), 1.55-1.80 (m, 6H), 1.81-1.90 (m, 1H), 1.90-2.06 (m, 1H), 2.07-2.20 (m, 2H), 2.20-2.30 (m, 2H), 2.85-2.97 (m, 2H), 3.00-3.07 (m, 3H), 3.19 (dd, *J* = 5.0; 10.0 Hz, 1H), 3.55 (d, *J* = 15.0 Hz, 2H), 3.73 (partially resolved dt, *J* = 2.5; 7.5 Hz, 2H estimated since accurate integration difficult due to broad water peak centered at 3.70), 3.98 (partially resolved dd, *J* = 5.0; 15.0 Hz, 1H) partial overlap with 4.04 (d, *J* = 15.0 Hz, 2H), 4.19-4.25 (m, 1H estimated), 4.41 (m, 1H), 4.50-4.56 (m, 1H), 6.81 (br s, 1 H), 7.15 (d, *J* = 5.0 Hz, 2H), 7.22 (t, *J* = 5.0 Hz, 2H), 7.29 (t, *J* = 7.5 Hz, 1H), 7.32-7.43 (m, 3H), 7.49 (br s, not integrated), 7.56 (s, 1H) partial overlap with 7.60 (br s, 2H), 8.09 (d, *J* = 10.0 Hz, 1H), 8.39 (br s, 2H), 8.56 (br s, 1H), 8.78 (d, *J* = 5.0 Hz, 1H), 8.99 (br s, 1H), 9.20 (br s, 1H)

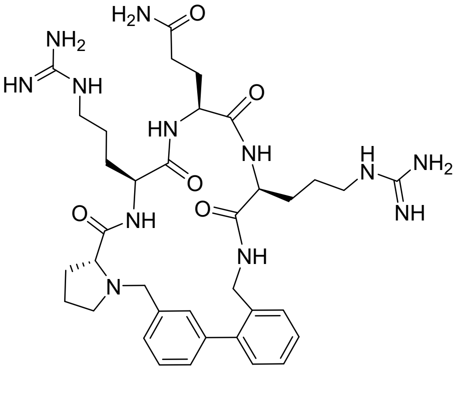

CY007455

LC-MS: $t_R$ 1.80 min, 100% (UV, 220 nm), [(M+H)$^+$] 733.50

UPLC: $t_R$ 2.42 min, 100% (UV, 220 nm)

**CY007458**

$^1$H NMR (DMSO-d$_6$, 700 MHz): δ 1.06-1.12 (m, 1H), 1.13-1.24 (m, 2H), 1.35-1.42 (m, 1H), 1.53-1.84 (m, 8H), 2.03-2.21 (m, 3H), 2.35 (br q, *J* = 7.0 Hz, 1H), 2.84-2.94 (m, 2 H), 3.05-3.14 (m, 3H), 3.21 (dd, *J* = 7.0; 7.0 Hz, 1H), 3.62 (d, *J* = 14.0 Hz, 2H), 3.93 (partially resolved q, *J* = 7.0 Hz, 1H), 3.97 (d, *J* = 14.0 Hz, 1H), 4.02-4.13 (m, 2H), 4.31 (br s, 1H), 4.41 (br q, *J* = 7.0 Hz, 1H), 6.81 (s, 1H), 7.14 (t, *J* = 7.0 Hz, 1H), 7.19 (d, *J* = 7.0 Hz, 1H), 7.21 (d, *J* = 14.0 Hz, 1H), 7.28 (t, *J* = 7.0 Hz, 1H), 7.34-7.38 (m, 2H), 7.44 (s, 1H), 7.54 (s, 1H), 7.61(br s, 3H), 7.75 (br s, 2H), 7.88 (t, *J* = 7.0 Hz, 1H), 8.39 (br s, 3H), 8.95 (d, *J* = 7.0 Hz, 1H), 8.98 (partially resolved d, *J* = 7.0 Hz, 2H)

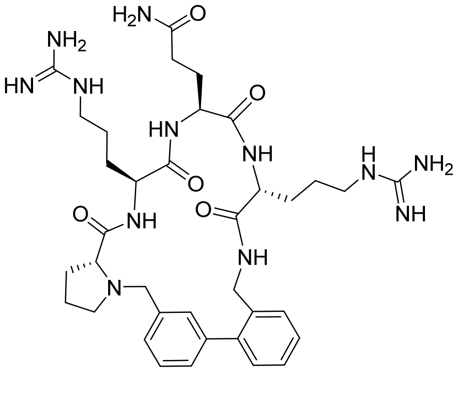

CY007458

$^{13}$C NMR (DMSO-d$_6$, 176 MHz): δ 23.59, 23.64, 25.4, 25.8, 27.9, 29.9, 30.97, 31.03, 31.3, 38.9, 40.37, 40.43, 49.7, 53.2, 53.9, 54.3, 58.5, 67.5, 126.46, 126.50, 126.9, 127.6, 127.7, 128.4, 129.5, 136.3, 139.7, 140.7, 141.0, 157.2, 157.4, 166.4, 171.8, 172.2, 172.3, 172.4, 173.1

LC-MS: $t_R$ 1.79 min, 100% (UV, 220 nm), [(M+H)$^+$] 733.50

HRMS (ESI-TOF, [M+H]$^+$, *m/z*): Calculated for $C_{36}H_{53}N_{12}O_5$ 733.42564, Found 733.42817

**CY007476**

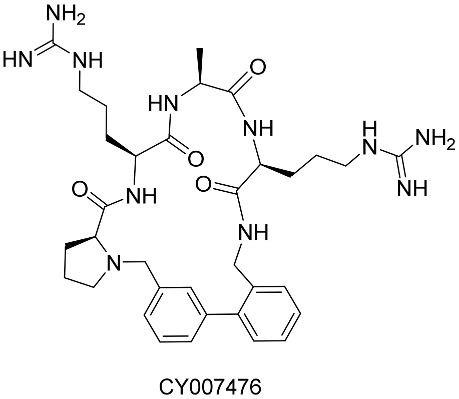

CY007476

$^{1}H$ NMR (DMSO-$d_6$, 700 MHz): δ 1.24 (d, $J$ = 14.0 Hz, 3H), 1.30-1.37 (m, 1H), 1.43-1.53 (m, 2H), 1.56-1.67 (m, 2H), 1.67-1.76 (m, 3H), 1.80 (br s, 1H), 1.84-1.91 (m, 1 H), 2.08-2.15 (m, 1H), 2.26 (q, $J$ = 7.0 Hz, 1H), 2.78 (br m) partial overlap with 2.82 (t, $J$ = 7.0 Hz, total 2H), 3.02 (br m) partial overlap with 3.05 (dd, $J$ = 7.0; 10.5 Hz, total 3H), 3.30 (d, $J$ = 14.0 Hz, 1H), 3.77-3.80 (dq, $J$ = 2.5; 7.0 Hz, 1H), 3.93-4.00 (m, 2H), 4.04 (d, $J$ = 7.0 Hz), 4.40-4.47 (m, 2H), 7.13 (d, $J$ = 7.0 Hz, 1H), 7.20 (d, $J$ = 7.0 Hz, 1H), 7.25-7.29 (m, 2H), 7.31 (t, $J$ = 7.0 Hz, 1.5H), 7.36-7.41 (m, 2.5H), 7.59 and 7.61 (overlapping br s, total 2H), 8.02 (d, $J$ = 7.0 Hz, 1H), 8.44 (br s, not integrated, multiple H), 8.55 (br s, 1H), 8.95 (v br s, 1H), 9.07 (v br s, 1H), 9.18 (v br s, 1H)

$^{13}C$ NMR (DMSO-$d_6$, 176 MHz): δ 16.8, 23.3, 24.2, 25.0, 27.2, 28.8, 29.8, 38.9, 40.3, 40.4, 50.5, 51.4, 52.7, 53.6, 58.7, 67.7, 126.8, 127.4, 127.6, 127.9, 128.5, 128.9, 129.7, 129.8, 137.0, 139.1, 140.6, 140.9, 157.6, 165.9, 172.1, 173.3, 173.5, 173.6

LC-MS: $t_R$ 1.97 min, 100% (UV, 220 nm), [$(M+H)^+$] 676.40

HRMS (ESI-TOF, $[M+H]^+$, $m/z$): Calculated for $C_{34}H_{50}N_{11}O_4$ 676.40418, Found 676.40702

**CY007491**

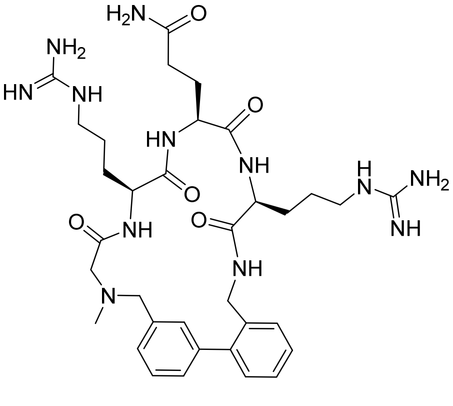

CY007491

$^{1}H$ NMR (DMSO-$d_6$, 700 MHz): δ 1.18-1.32 (m, 2H), 1.43-1.52 (m, 2H), 1.60-1.68 (m, 2H), 1.85-1.97 (m, 2H), 2.08-2.15 (m, 1H), 2.18-2.25 (m) partial overlap with 2.19 (s, total 4H), 2.88 (br s, 2H), 3.03-3.18 (m, 4H), 3.52 (d, $J$ = 14.0 Hz, 1H), 3.63 (d, $J$ = 14.0 Hz, 1H), 3.71-3.76 (m, 1H), 4.09 (br d, $J$ = 14.0 Hz, 1H), 4.14-4.19 (br m, 1H), 4.20-4.26 (m, 1H), 4.46 (dt, $J$ = 3.5, 7.0 Hz, 1H), 6.79 (br s, 2H), 7.16 (dd, $J$ = 7.0; 14.0 Hz, 2H), 7.24 (overlapped dd, $J$ = 7.0; 7.0 Hz, 2H), 7.28 (t, $J$ = 7.0 Hz, 1H), 7.34 (t, $J$ = 7.0 Hz, 1H), 7.40 (t, $J$ = 7.0 Hz, 1H), 7.46 (s, 1H), 7.54 (s, 1H), 7.91 (v br s, not integrated), 8.02 (d, $J$ = 14.0 Hz, 1H), 8.43 (br s, not integrated), 8.63 (d, $J$ = 7.0 Hz, 1H), 8.85 (br s, 2H), 9.09 (v br s, 2H)

$^{13}C$ NMR (DMSO-$d_6$, 176 MHz): δ 23.2, 25.2, 26.2, 27.0, 29.6, 31.6, 39.0, 40.3, 40.4, 42.2, 42.4, 50.5, 52.9, 56.1, 60.9, 61.1, 126.7, 127.4, 127.6, 128.0, 128.2, 128.5, 129.6, 136.3, 138.8, 140.5, 141.0, 157.5, 157.6, 162.9, 165.8, 169.5, 171.6, 172.9, 173.5

LC-MS: $t_R$ 1.86 min, 100% (UV, 220 nm), [$(M+H)^+$] 707.30

HRMS (ESI-TOF, $[M+H]^+$, *m/z*): Calculated for $C_{34}H_{51}N_{12}O_5$ 707.40999, Found 707.41313

**CY007493**

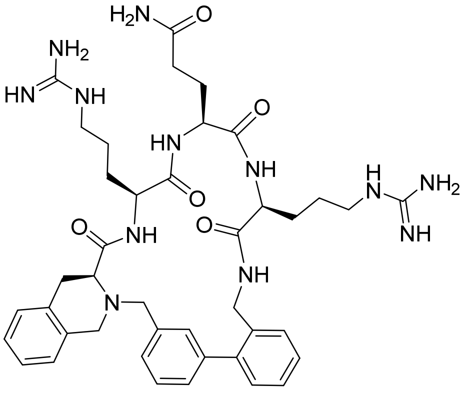

CY007493

$^1H$ NMR (DMSO-$d_6$, 700 MHz): δ 1.13 (br s, 1H), 1.23 (m, 2H), 1.43-1.67 (m in three parts, 5H), 1.83-1.98 (m in two parts, 3H), 2.08-2.26 (m in 2 parts, 3H), 2.83 (br s, 2H), 2.95-3.07 (m, 5 H), 3.60 (br d, *J* = 14.0 Hz, 2H estimated due to very broad and large water peak making accurate integration difficult in 3.50-4.00 region), 3.70-3.75 (br m, 2H estimated), 3.79 (br s, not integrated), 3.83 (br s, not integrated), 4.10 (v br s, not integrated), 4.15 (br s, 1H), 4.25 (v br s, not integrated), 4.42 (br s, 1H), 6.80 (br s, 1H), 7.04 (d, *J* = 7.0 Hz, 1H), 7.13 (t, *J* = 7.0 Hz, 1H) partial overlap with 7.15-7.20 (m, 4H), 7.27 (br d, *J* = 14.0 Hz, 1H) partial overlap with 7.30 (t, *J* = 7.0 Hz, 1H), 7.35 (d, *J* = 7.0 Hz, 1H), 7.37 (t, *J* = 7.0 Hz, 1H), 7.43 (br s, 1H), 7.61 (v br s, not integrated) partial overlap with 7.66 (br s, 1H), 8.04 (v br s, 2H), 8.42 (br s, 3H), 8.63 (br s, 3H), 8.96 (v br s, 2H)

$^{13}C$ NMR (DMSO-$d_6$, 176 MHz) δ 22.1, 23.2, 24.5, 25.2, 26.2, 26.7, 28.0, 29.1, 29.5, 31.3, 31.6, 38.9, 40.36, 40.43, 52.1, 52.9, 56.0, 61.9, 126.2, 126.4, 126.7, 126.8, 127.2, 127.6, 127.9, 128.4, 128.7, 129.5, 133.6, 133.7, 136.4, 138.7, 140.8, 141.1, 157.2, 157.3, 167.0, 171.5, 172.9, 173.5

LC-MS: $t_R$ 2.13 min, 100% (UV, 220 nm), [$(M+H)^+$] 795.50

HRMS (ESI-TOF, $[M+Na+H]^{+2}$, *m/z*): Calculated for $C_{41}H_{55}N_{12}O_5(Na)$ 409.21526, but since doubly charged 818.43052, Found 409.22007 [(x 2) 818.44014]

**CY007502**

$^{1}$H NMR (DMSO-$d_6$, 500 MHz): δ 1.28-1.46 (br m in two parts, 4H), 1.50-1.62 (m, 2H), 1.71-1.97 (m in at least two parts, 4H), 2.06-2.37 (m in three parts, 4H), 2.81-3.06 (m in two parts, 3H), 3.14-3.20 (m, 1H), 3.42 (d, *J* = 10.0 Hz, 1H), 3.65-3.68 (partially resolved dt, *J* = 2.5; 7.5 Hz, 1H), 3.98.-4.04 (m, approximates t, 2H), 4.20-4.31 (br m, 2H), 4.36-4.54 (br m, 4H), 6.72 (br d, *J* = 10.0 Hz, 1H), 6.78 (br s, 1H), 7.11-7.19 (m, 2H), 7.29 (dt, *J* = 2.5, 5.0 Hz, 1H), 7.35 (partially resolved dq, *J* = 2.5; 5.0 Hz, 2H), 7.44 (br s, 1H), 7.59 (partially resolved dd, *J* = 2.5; 10.0 Hz, 2H), 7.84 (v br s, not integrated), 8.12 (d, *J* = 10.0 Hz, 1H), 8.36 (br, d, *J* = 10.0 Hz) overlaps with 8.43 (br s, total 5H), 8.82 (br s, 1H), 9.14 (m, 1H)

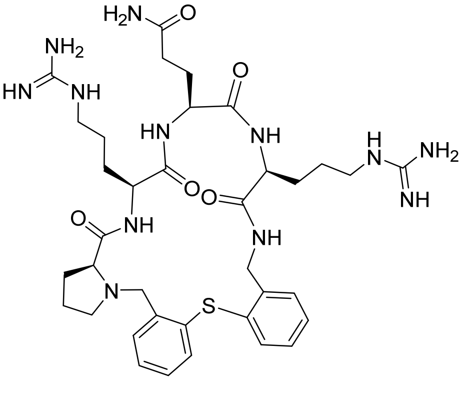

CY007502

LC-MS: $t_R$ 2.03 min, 100% (UV, 220 nm), [(M+H)$^+$] 765.50

UPLC: $t_R$ 2.73 min, 100% (UV, 220 nm)

**Supplementary Information References**

[16] Wahhab, A., Thomas, H., Richard, L., Peterson, M. L., Macdonald, D. & Dubé, D. Libraries of heteroaryl-containing macrocyclic compounds and methods of making and using the same. *Intl. Pat. Publ. No. WO 2017/049383*, filed 14 September 2016.

[17] Macdonald, D., Dubé, D.; Wahhab, A., Thomas, H., Richard, L. & Peterson, M. L. Libraries of diverse macrocyclic compounds and methods of making and using the same. *Intl. Pat. Publ. No. WO 2017/197488*, filed 16 May 2017.

[18] Wahhab, A.; Dubé, D.; Macdonald, D.; Peterson, M. L.; Richard, L. & Thomas, H. Libraries of pyridine-containing macrocyclic compounds and methods of making and using the same. *Intl. Pat. Publ. No. WO 2018/232506*, filed 20 June 2018.

[19] Kaiser, E., Colescott, R. L., Bossinger, C. D. & Cook, P. I. Color test for detection of free terminal amino groups in the solid-phase synthesis of peptides. *Anal. Biochem.* **34,** 595-598 (1970).

[20] Similarly to has been done using the 2-hydroxy-4-methoxybenzyl (Hmb) for protection of the backbone amide bond: Zeng, W., Regamey, P. O., Rose, K., Wang, Y. & Bayer, E. Use of Fmoc-N-(2-hydroxy-4-methoxybenzyl)amino acids in peptide synthesis. *J. Pept. Res.* **49,** 273-279 (1997).

[21] Fraser, G., Marsault, E., Peterson, M., Hoveyda, H., Beaubien, S., Benakil, K. & Deziel, R. Macrocyclic antagonists of the motilin receptor. *Intl. Pat. Publ. No. WO 2004/111077*, filed 18 June 2004.
